# Supervised Application of Internal Validation Measures to Benchmark Dimensionality Reduction Methods in scRNA-seq Data

**DOI:** 10.1101/2020.10.29.361451

**Authors:** Forrest C Koch, Gavin J Sutton, Irina Voineagu, Fatemeh Vafaee

## Abstract

A typical single-cell RNA sequencing (scRNA-seq) experiment will measure on the order of 20,000 transcripts and thousands, if not millions, of cells. The high dimensionality of such data presents serious complications for traditional data analysis methods and, as such, methods to reduce dimensionality play an integral role in many analysis pipelines. However, few studies benchmark the performance of these methods on scRNA-seq data, with existing comparisons assessing performance via downstream analysis accuracy measures which may confound the interpretation of their results. Here, we present the most comprehensive benchmark of dimensionality reduction methods in scRNA-seq data to date, utilizing over 300,000 compute hours to assess the performance of over 25,000 low dimension embeddings across 33 dimensionality reduction methods and 55 scRNA-seq datasets (ranging from 66-27,500 cells). We employ a simple-yet-novel approach which does not rely on the results of downstream analyses. Internal validation measures (IVMs), traditionally used as an unsupervised method to assess clustering performance, are repurposed to measure how well-formed biological clusters are after dimensionality reduction. Performance was further evaluated using nearly 200,000,000 iterations of DBSCAN, a density-based clustering algorithm, showing that hyperparameter optimization using IVMs as the objective function leads to near-optimal clustering. Methods were also assessed on the extent to which they preserve the global structure of the data, and on their computational memory and time requirements across a large range of sample sizes. Our comprehensive benchmarking analysis provides a valuable resource for researchers and aims to guide best practice for dimensionality reduction in scRNA-seq analyses, and we highlight LDA (Latent Dirichlet Allocation) and PHATE (Potential of Heat-diffusion for Affinity-based Transition Embedding) as high-performing algorithms.

---

Single-cell RNA sequencing (scRNA-seq) is revolutionizing our understanding of gene expression, from cataloguing cell-types^1^ to elucidating transcriptional regulation^2–4^. Consequently, recent years have seen a drastic increase in the number of software packages devoted to the analysis of scRNA-seq data. At the time of writing, scRNA-tools.org tracks over 500 such packages, up from around 100 in 2017^5^. Furthermore, recent research has suggested that many software packages originally intended for the analysis of bulk RNA-seq data may be just as performant, if not more so, than their scRNA-seq devoted counterparts^6^. As a result, efforts to benchmark these software packages have become vital resources for researchers who wish to determine the best way to analyze their data. Benchmarking efforts are further complicated by the fact that many of these packages are intended to perform multiple stages in the analysis of scRNA-seq data^7–9^, and comparisons of complete pipelines may fail to answer the question of why one pipeline may outperform another. To answer this question, we must consider the various components of scRNA-seq analysis separately.

Dimensionality reduction—the process of extracting a low dimensional structure from a higher dimensional set of data—is an essential component of scRNA-seq analysis. It is used to extract important biological conclusions from scRNA-seq data, including the identification of novel cell-types and disease-specific gene expression changes. Therefore, comprehensive benchmarking of the accuracy of dimensionality reduction methods is particularly important.

Dimensionality reduction is used extensively in a wide range of research from signal and image processing to epidemiology^10,11^. The widespread usage of dimensionality reduction can be largely attributed to its ability to mitigate the negative effects of the so-called “curse of dimensionality” ^12^.

In 2019, Sun *et al*. benchmarked 18 dimension reduction methods against 30 datasets^13^. However, results were largely dataset dependent, and sample size was a limiting factor in this study with only two datasets exceeding 1,000 cells. In 2020, Tsuyuyaki *et al*. compared 21 different implementations of Principal Component Analysis (PCA) over 22 datasets^14^. However, common PCA algorithms such as kernel PCA were not included. Also, performance in this study was assessed by agreement with other the methods, which may result in the most “average” PCA implementation receiving the best results. Furthermore, both studies assessed performance after downstream clustering analysis. As previously mentioned, such an approach is limited as they may be confounded by the downstream analysis. Recent work by Heiser and Lau (2020) has attempted to address this limitation by considering the ability of a dimensionality reduction method to preserve structures in the data^15^, however, such an approach fails to consider separability of existing clusters in the data and may be biased towards linear methods such as PCA.

In this work, we present the most comprehensive benchmark of dimensionality reduction methods in scRNA-seq data to date. Utilizing over 300,000 compute hours, we assessed the performance of 24,824 low dimension embeddings, representing 33 methods, 55 datasets, 20 tissue types, and 12 sequencing protocols. These datasets ranged from 66–27,500 cells and contained between 3 and 56 distinct cell-types.

We employed a novel approach to comparing dimensionality reduction methods which does not rely on downstream analyses such as clustering accuracy. To achieve this, we propose the *supervised* application of internal validation measures (IVMs) to quantify how compact and well-separated biological clusters appear after dimensionality reduction. These measures have traditionally been used as an *unsupervised* approach to determine the ideal clustering of a group of data (*i.e*., selecting the optimal value of k during k-means clustering)^16^. Using “ground-truth” labels to calculate an IVM allows for the quantification of how well-defined the existing biological clusters appear in the data. This enabled the comparison of dimensionality reduction methods based on their tendency to produce low dimension embeddings with well-defined biological clusters.

We then explored whether these results translate into higher accuracies during clustering by analyzing the results of nearly 200,000,000 iterations of DBSCAN, a commonly used densitybased clustering algorithm^17^. Hyperparameter optimization was performed using IVMs as the objective function, and the resulting accuracy was assessed for each method. Additionally, we consider how well embeddings, chosen based on these measures, preserve the global structure of the data, as well as their memory and time requirements. This benchmark covers a wider range of dimensionality reduction methods than any scRNA-seq benchmark to date and attempts to provide method developers with a comprehensive resource summarizing the properties of a wide range of dimensionality reduction methods.

## Results

We assessed the performance of 33 dimensionality reduction methods (Table 1) across 55 publicly available scRNA-seq datasets (Supplementary Table 1). These data sets cover a wide range of sequencing techniques that including Smart-seq2 (22 datasets), Chromium (10), SMARTer (7), STRT-seq (4), Drop-seq (3), inDrop (3), C1 (3), and others (3). Additionally, selected list of datasets covers 24 different tissue types with diverse number of cells (66 – 27,499), cell types (3 – 56).

**Table 1.** Dimensionality methods that were benchmarked

| Method (abbreviation) | Brief Description | Linear/Non-Linear | Package |
| --- | --- | --- | --- |
| Bayesian decomposition (bd)<sup>28</sup> | Uses a Gibbs sampler to estimate the posterior density of NMF factors | Linear | nimfa<sup>29</sup> |
| Factor Analysis (fa)<sup>30</sup> | Linear generative model with Gaussian latent variables | Linear | sklearn<sup>25</sup> |
| Fast Independent Component Analysis (fica)<sup>31</sup> | Extracts statistically independent non-Gaussian signals | Linear | sklearn |
| Gaussian Random Projection (grp)<sup>32</sup> | Projecting the original input space on a randomly generated matrix where components are drawn from a normal distribution | Linear | sklearn |
| Incremental PCA (ipca)<sup>33</sup> | A mini-batch implementation of PCA | Linear | sklearn |
| Isometric Mapping (isomap)<sup>34</sup> | An extension on MDS that attempts to preserve geodesic distances | Non-linear | sklearn |
| Iterated Conditional Modes (icm)<sup>28</sup> | Similar to BD, but uses modes in place of random samples in the Gibbs sampler | Linear | nimfa |
| ivis<sup>35</sup> | Uses Siamese neural networks to embed data | Non-linear | ivis |
| Kernel PCA (kPCA)<sup>36</sup> (with cosine (cos), polynomial (pol), and radial basis function (rbf) transformations) | Utilizes the kernel trick to allow for non-linear representations of the data. | Non-linear | sklearn |
| Latent Dirichlet allocation (lda)<sup>37</sup> | Bayesian generative model utilizing a Dirichlet prior over latent space | Non-linear | sklearn |
| Least squares NMF (lsnmf)<sup>38</sup> | Alternative to multiplicative NMF which claims to have faster convergence | Linear | nimfa |
| Locally linear embedding (lle)<sup>39</sup> | Similar to isomap, but attempts to preserve local distances | Non-linear | sklearn |
| Non-negative Matrix Factorization (nmf-nnsvd and nmf-lee)<sup>40,41</sup> | Factorizes a (non-negative) data into two non-negative matrices. | Linear | sklearn/nimfa |
| Non-smooth NMF (nsnmf)<sup>42</sup> | Applies smoothing kernel into the factorization step reducing sparsity | Linear | nimfa |
| Potential of Heat-diffusion for Affinity-based Trajectory Embedding (phate)<sup>43</sup> | Utilizes a heat diffusion model to preserve branching structures in data | Non-linear | phate |
| Principle Component Analysis (pca)<sup>44,45</sup> | Construct a low dimensional, orthogonal basis that maximally explains observed variance in the data. | Linear | sklearn |
| Probabilistic non-negative Matrix Factorization (pmf)<sup>46</sup> | Assumption that data follows a multinomial distribution and performs expectation maximization | Linear | nimfa |
| Probabilistic sparse MF (psmf)<sup>47</sup> | Assumption of Gaussian noise within the data, and sparsity constraint added for factorization | Linear | nimfa |
| Sparse Autoencoder for Clustering, Imputing, and Embedding (saucie)<sup>48</sup> | Sparse Autoencoder Neural Network | Non-linear | saucie |
| Sparse nmf (snmf)<sup>49</sup> | Addition of sparsity constraint to factorization process | Linear | nimfa |
| sparse PCA (spca and spca-batch)<sup>50</sup> | An extension of PCA which aims to produce sparse (with many zeros) components | Linear | sklearn |
| Sparse Random Projection (srp)<sup>51</sup> | Similar to GRP, but matrix elements are drawn from a zero-inflated, centered uniform distribution. | Linear | sklearn |
| Laplacian eigenmaps (spectral)<sup>52,53</sup> | Attempts to reconstruct the manifold in low dimensional space by preserving the distances of the nearest neighbours to each point | Non-linear | sklearn |
| t-distributed Stochastic Neighbour Embedding (t-SNE)<sup>54</sup> | Minimizes the KL divergence of the estimated distribution of points in high dimensional points with a similar low-dimensional distribution. | Non-linear | Mutlicore TSNE<sup>55</sup> |
| Truncated Singular Value Decomposition (tsvd)<sup>44</sup> | PCA without centring and scaling | Linear | sklearn |
| Uniform Manifold Approximation and Projection (umap)<sup>56</sup> | Constructs a fuzzy topological representation of the data to preserve in the low dimensional embedding | Non-linear | Umap-learn |
| Variational autoencoder (vasc)<sup>31</sup> | Uses neural network to map latent variables to zero-inflated negative binomial distribution | Non-linear | vasc |
| Variational projection (vpac)<sup>57</sup> | Uses a variational projection algorithm to infer a Gaussian mixture in latent space | Non-linear | vpac |
| Zero-inflated Factor Analysis (zifa)<sup>58</sup> | Uses a latent variable model, similar to FA, in which is modulated by the addition of a zero-inflated layer meant to simulate dropout. | Linear | ZIFA |

Both counts-per-million (CPM) and log-transformed CPM data for each dataset were used, and embeddings varied between 2 and 96 dimensions were generated (Fig. 1). Unless otherwise stated, when applying each method to each dataset, only the best performing embedding and transformation combination was reported. In accordance with the benchmark guidelines set out by Weber *et al*. 2019, no method was given special treatment^18^. Default arguments were used for all methods where applicable.

**Fig. 1:**
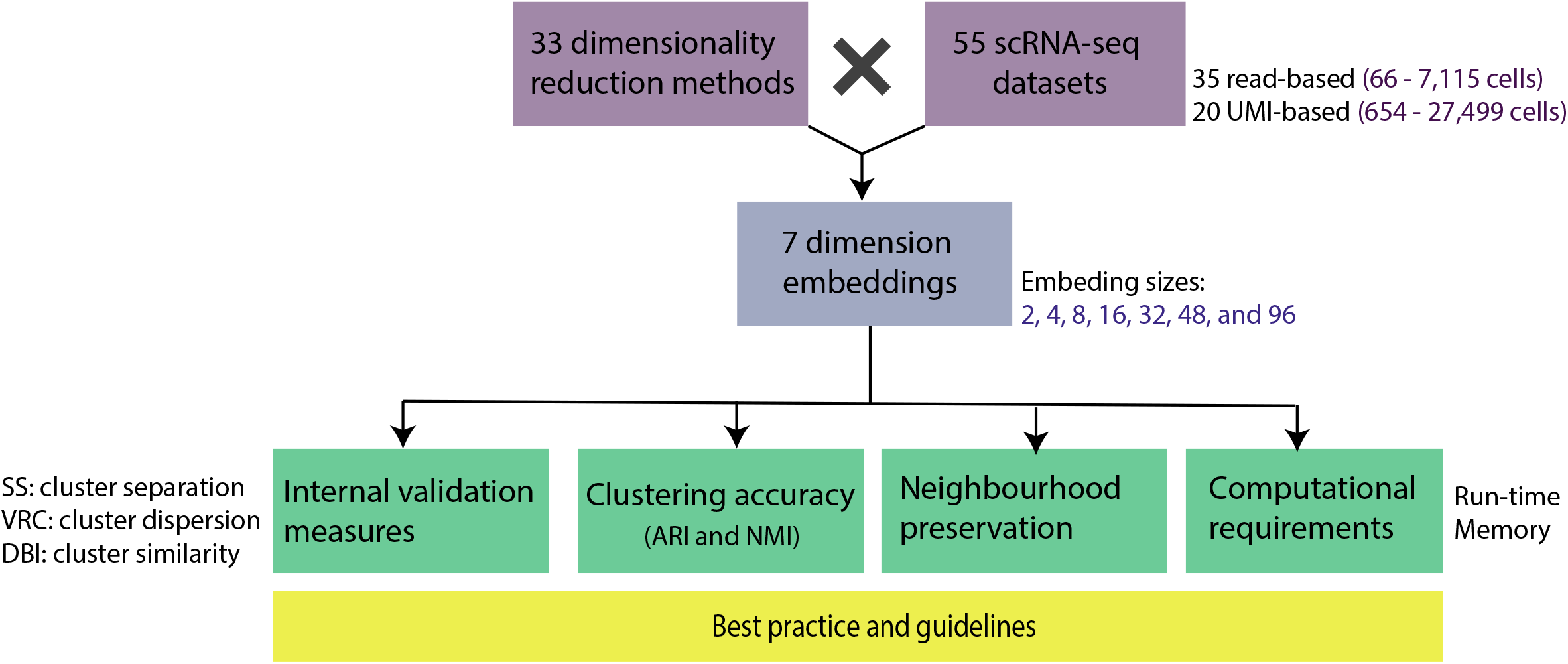
Overview of dimensionality reduction methods and benchmarking workflow. **(a)** Each of the 33 dimensionality reduction methods was applied to each of the 55 datasets. For each dataset, embeddings of dimension 2, 4, 8, 16, 32, 48, and 96 were obtained for both log1p-transformed and non-transformed data after count-per-million normalization, resulting in the total of 25,410 embeddings for this benchmarking. IVM, global structure, clustering accuracy, and resources analyses were performed.

### Supervised application of internal validation measures (IVMs)

To assess how well the different dimensionality reduction methods capture biologically-relevant clusters, we calculated internal validation measures (IVMs) for groups of cells defined by cell-type annotations provided with each published dataset. We used three IVMs commonly used for cluster validation^16,19^: silhouette score (SS)^20^, variance ratio criterion (VRC)^21^, and Davies Bouldin index (DBI)^22^.

A summary heatmap of IVMs across all datasets and methods is available in Fig. 2a. Dimensionality reduction methods were ranked according to the three IVMs on each dataset. Rank-based analyses were performed (see Methods for details) in order to make minimal assumptions about the distribution of each IVM per dataset, as well as to allow for comparisons between IVMs.

**Fig. 2:**
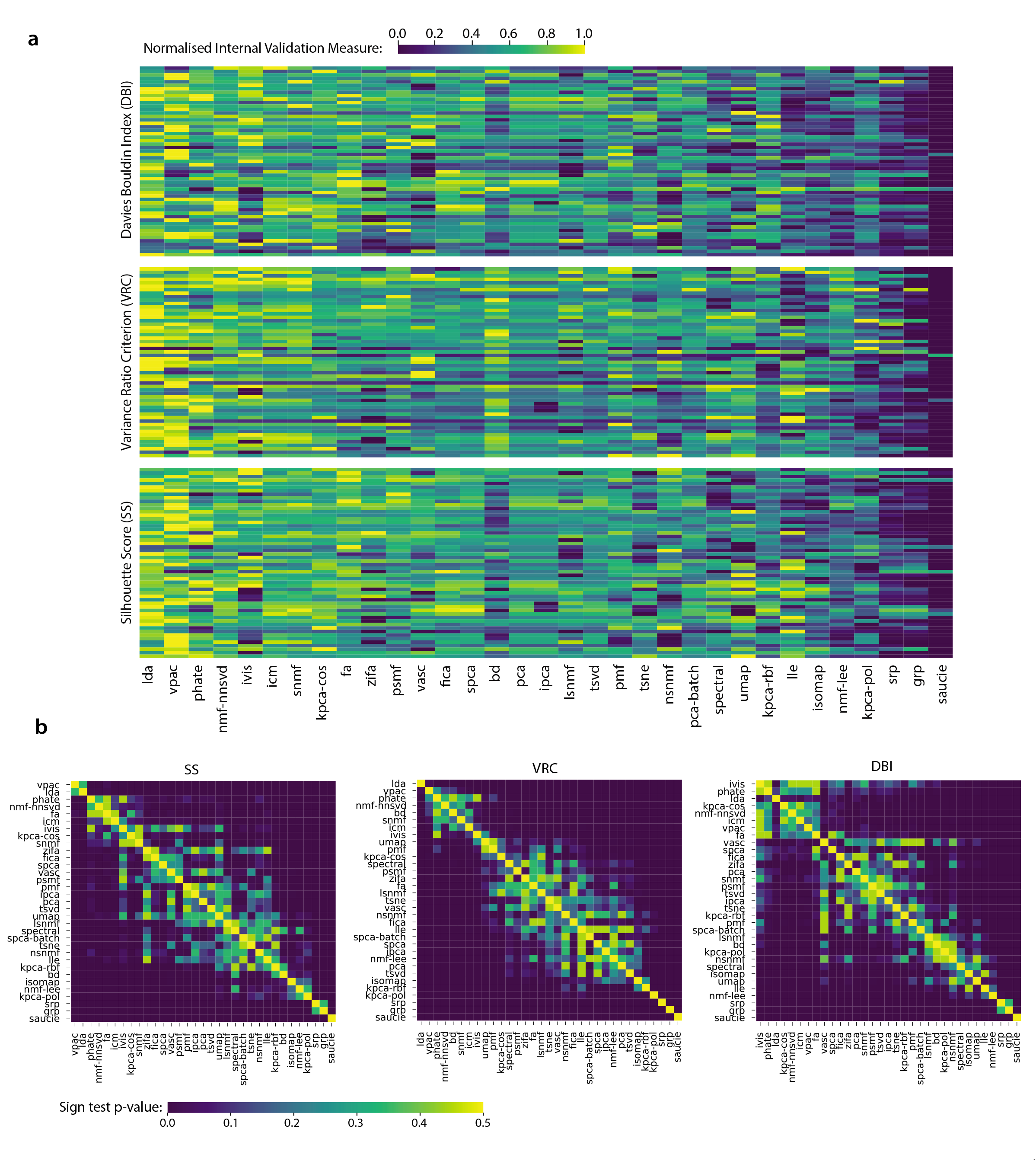
Overview of Internal Validation Measure (IVM) analysis. **(a)** Heatmap of the 3 IVMs (SS with Euclidean distance, VRC and DBI) for each dimensionality reduction method (columns). Values were standardized using min-max normalization applied per-dataset (rows) to highlight differences between methods. SS measures with three other distance metrics (cosine, correlation, and standardized Euclidean) are shown in Supplementary Fig. 1. **(b)** Heatmaps of p-values from pairwise sign-tests comparing IVMs for each method across all datasets. It depicts the relative ranking of dimensionality reduction methods, as well as providing information as to which methods were roughly equivalent.

LDA (average median rank = 4.0), VPAC (5.33), PHATE (6.0), and NMF (7.0), had the best average median ranks across all IVMs (Supplementary Table 2), however pairwise sign tests indicated that only LDA significantly outperformed all other methods (Fig. 2b, uncorrected p < 0.05). Interestingly, clusters of roughly equivalent performance were observed. For example, when optimizing embeddings for VRC, we observed no significant difference between the 2^nd^-8^th^ best methods, however, they significantly outperformed the remaining methods.

Despite their recent popularity for dataset visualization, UMAP (average median rank of 19.67) and t-SNE (17.33) performed relatively poorly. The widely used PCA exhibited similar performance (16.33).

We observed high concordance in the rankings of the dimensionality reduction methods between each of the IVMs (Kendall’s W = 0.83, p < 0.001). Furthermore, to assess the effect of the distance metrics on silhouette scores, we considered Euclidean, standardized Euclidean, correlation, and cosine similarity measures and observed a high concordance between them (Methods; Kendall’s W = 0.91, p < 0.001) (Supplementary Fig. 1). In contrast, concordance between datasets was moderate regardless of the IVM considered (Methods; Kendall’s W = 0.47-0.52, p < 0.001).

We next explored how the scRNA-seq data properties contribute to method performance. First, given the increasing appreciation that scRNA-seq data generated with protocols that employ Unique Molecular Identifiers (UMIs) have different properties from read-based data^23^, we compared method performance between UMI-based datasets (n=20) and read-based datasets (n=35). The average median rank of each method was strongly correlated across the two datatypes (r=0.72; Supplementary Fig. 2). However, we note that UMAP’s relative performance was greater in UMI- than read-based datasets, with it even ranking as the best performing method in the former group.

Second, given that cell-type annotations were obtained from the original publications of scRNA-seq datasets, we investigated whether the performance of dimensionality reduction methods was affected by how these annotations were determined: (a) based on biological properties only, independently of downstream analyses of scRNA-seq data; (b) by clustering of expression data, or (c) by clustering after dimensionality reduction (Supplementary Table 1). We found that VRC showed some difference across the three categories of datasets in 9 of the 33 methods (p<0.05 uncorrected, ANOVA); however, none of these differences survived correction for multiple comparisons (Supplementary Fig. 3). These data suggest that IVMs are robust to the choice of cell-type classification method for the most performant dimensionality reduction methods.

Finally, we explored how log transformation of the expression data affects method performance. While for most methods there was overall good agreement between VRC obtained on log-transformed and non-transformed data, we noted that the log transformation strongly affected the outcome of kPCA-rbf, LSNMF and to a lesser extent kPCA-pol (Supplementary Fig. 4).

### Clustering accuracy after dimensionality reduction

We next conducted a cluster-based assessment of accuracy after dimensionality reduction. For each embedding previously computed, we defined clusters using the DBSCAN algorithm^17^, a density-based clustering algorithm which is able to recognize arbitrary shaped clusters. We performed a random search over the hyper-parameter space to estimate the optimal clustering with respect to each of the three IVMs and four distance measures (see Methods for further details). This optimization was unsupervised and did not make use of celltype labels. Adjusted Rand Index (ARI) was then used to calculate accuracy for each optimized embedding (Fig. 3a). Similar results were observed when using Normalized Mutual Information (NMI) to assess accuracy (Supplementary Fig. 5).

**Fig. 3:**
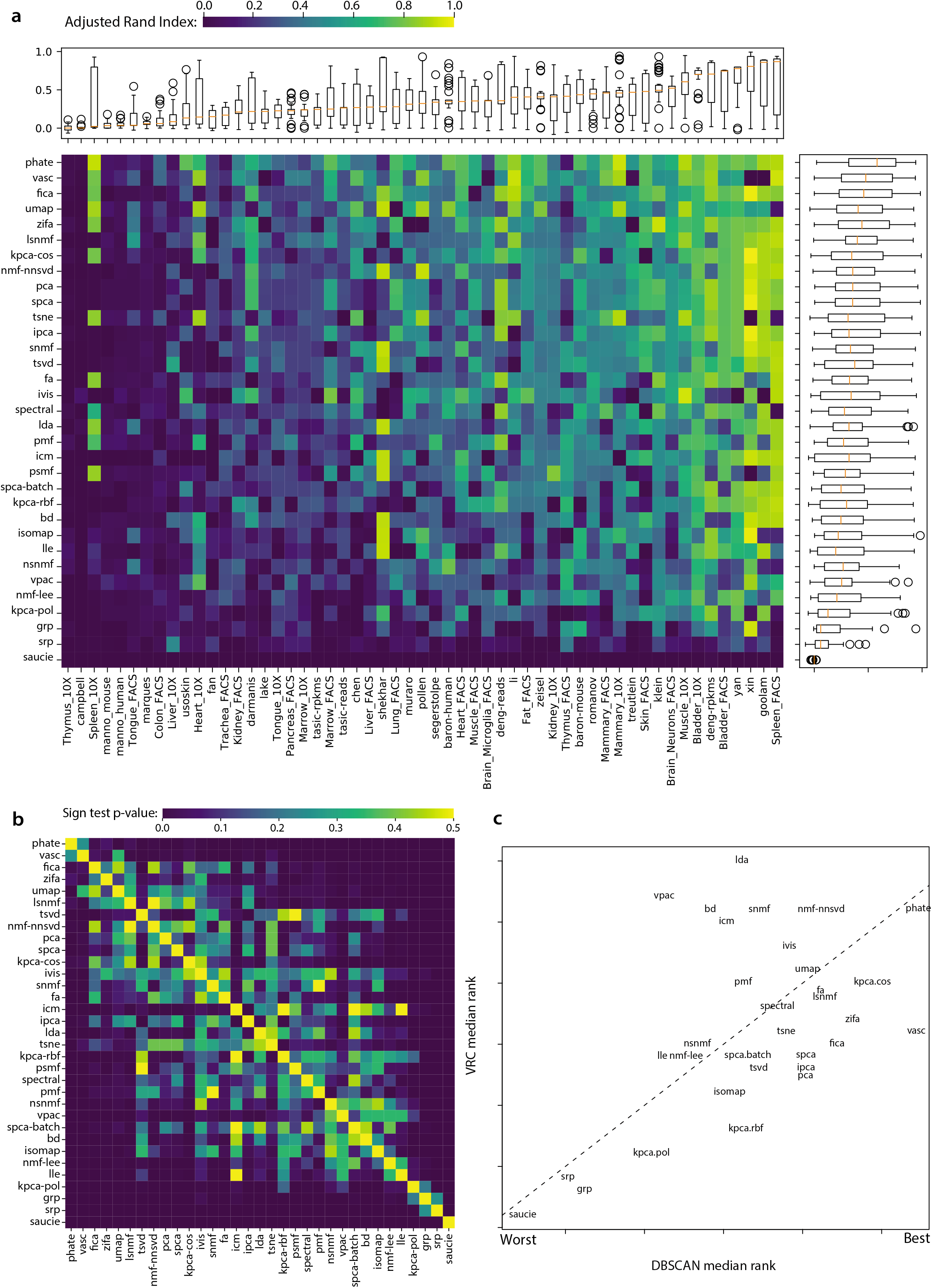
Overview of DBSCAN clustering analysis. **(a)** Heatmap with boxplots of the Adjusted Rand Index (ARI) achieved by each method (rows) for each dataset (columns). **(b)** Heatmaps of p-values from pairwise sign-tests comparing the accuracy of each method. Similar to Fig. 2b, it depicts the relative ranking of dimensionality reduction methods, as well as providing information as to which methods were roughly equivalent. **(c)** Comparison of the relative rankings assigned to each method by the IVM and DBSCAN clustering analyses. Best ranked methods appear in the top-right whereas worse ranked methods appear in the bottom left. The dotted line represents parity.

Supplementary Table 3 summarizes the average ARI score across datasets for each dimensionality reduction method, IVM (DBI, SS, VRC), and distance measure (Euclidean, standardized Euclidean, correlation, and cosine) combination. Most dimensionality reduction methods yielded their highest average ARIs when using VRC as the objective function with standardized Euclidean (SEU) distance metric for DBSCAN hyperparameter optimization (p < 0.001 by Monte Carlo simulation, see Methods).

PHATE (ARI = 0.53), VASC (0.46), FICA (0.45), and UMAP (0.44) resulted in the best average clustering accuracies (Supplementary Table 2). In general, clustering accuracy was highly dataset dependent (Fig. 3a). Although both Wilcoxon Sign-Rank test and Sign tests indicated that PHATE significantly outperformed most other methods (Fig. 3b), it is difficult to draw a conclusion regarding the ordering of the remaining methods. Overall, results in the clustering analysis were much noisier than the IVM analysis.

Nonetheless, to assess the agreement between these clustering analyses and the IVM analyses, the median rank across datasets was calculated for each method. A moderately strong linear (r=0.51, p=0.003; W=0.68, p=0.02) relationship was found between the two analyses (Fig. 3c).

### Effect of dimensionality reduction on global structure

To quantify how well global structure is preserved by these methods, we calculated pairwise distances between every cell before and after dimensionality reduction and compared them using a Spearman correlation. A high correlation indicates that the relative positions of each data point (*i.e*. cell) have not changed after dimensionality reduction thus preserving the structure of the data. Results for Euclidean distances are reported in-text, but other distance metrics were also considered and are reported in Supplementary Fig. 6. We selected embeddings with optimal DBI, SS, and VRC from the earlier IVM analysis to avoid considering dimensionality as a factor. Fig. 4 shows boxplots of correlations across all datasets.

**Fig. 4:**
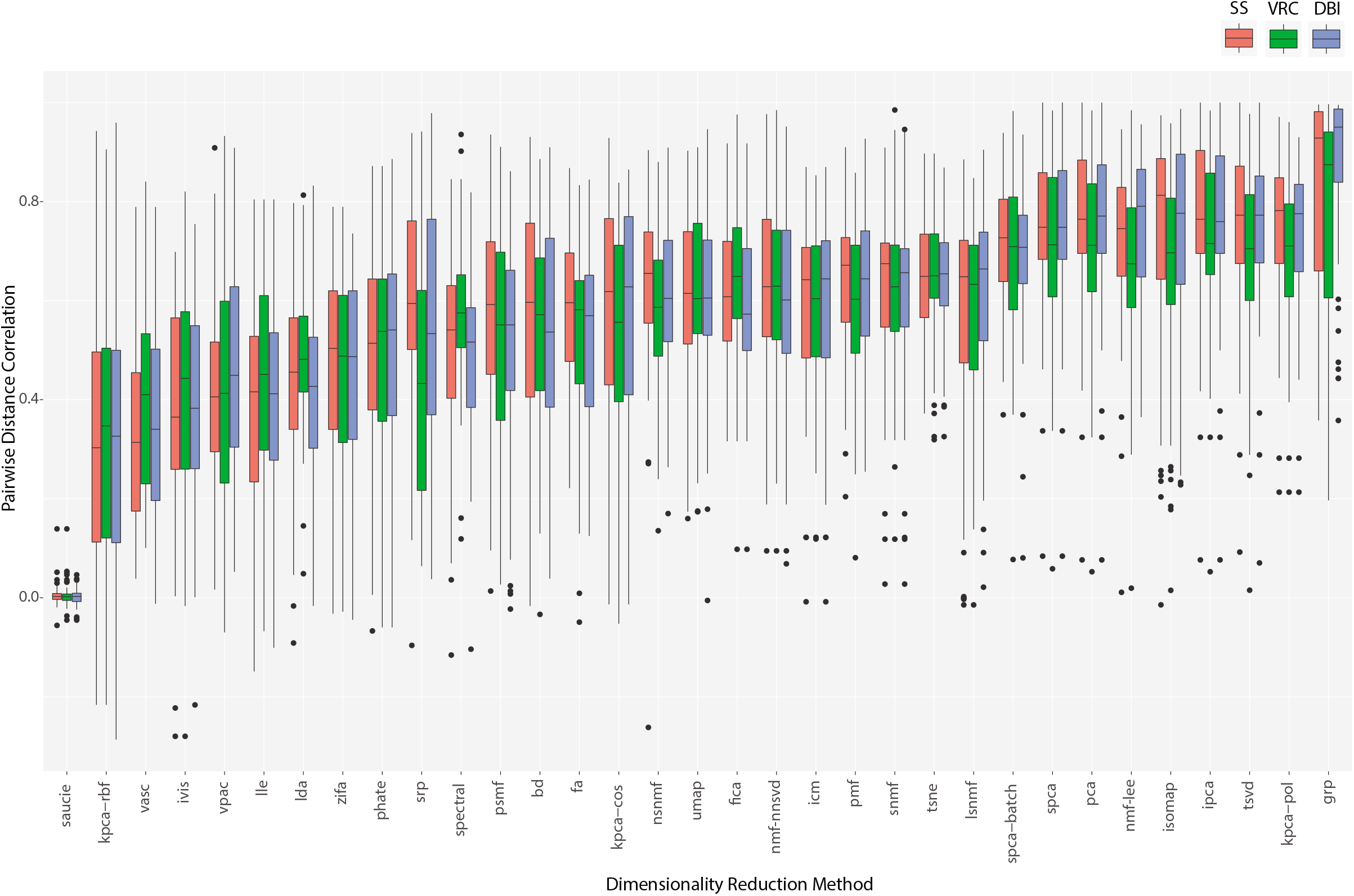
Preservation of global neighboring structure after dimensionality reduction. It shows boxplots of the spearman correlation between the pairwise distance matrices before and after dimensionality reduction for each dimensionality reduction method chosen according to the internal validation measure analysis (*c.f*. Fig. 1). Each point represents one dataset.

Of all algorithms tested, we found that GRP best preserved pairwise distances followed by TSVD and various PCA algorithms (Fig. 4). PCA’s performance in this assessment is consistent with results from Heiser and Lau (2020), who also found that PCA preserved local neighborhoods and global structure^15^. Interestingly, many of the methods that performed well in IVM and clustering analyses, such as PHATE, LDA, and VPAC, were ranked poorly here, and vice versa (*e.g*. GRP). This might suggest that, in single-cell data, the preservation of global structure is a weak indicator of a dimensionality reduction algorithm’s utility in detecting biological clusters.

We note that similar results were obtained using other distance measures for SS (W=0.90, p < 0.001), and using other IVMs (DBI, SS, VRC) to select the optimal embedding number (W=0.97, p < 0.001) (Supplementary Fig. 6).

### Qualitative comparisons of dimensionality reduction methods

In addition to quantitative comparisons, dimensionality reduction methods can also be compared visually through 2D projections. Fig. 5 presents 2-dimension embeddings of the 4 best performing methods according to clustering analyses for a few select datasets which were accurately clustered. We see that PHATE, VASC, and UMAP successfully separate celltypes into well-defined clusters despite being fully unsupervised methods. FICA, while keeping cells of the same type close together, did not produce as much separation between clusters. Furthermore, UMAP displayed a tendency to “fracture” the data, giving the impression of a structure in the data which may not actually exist.

**Fig. 5:**
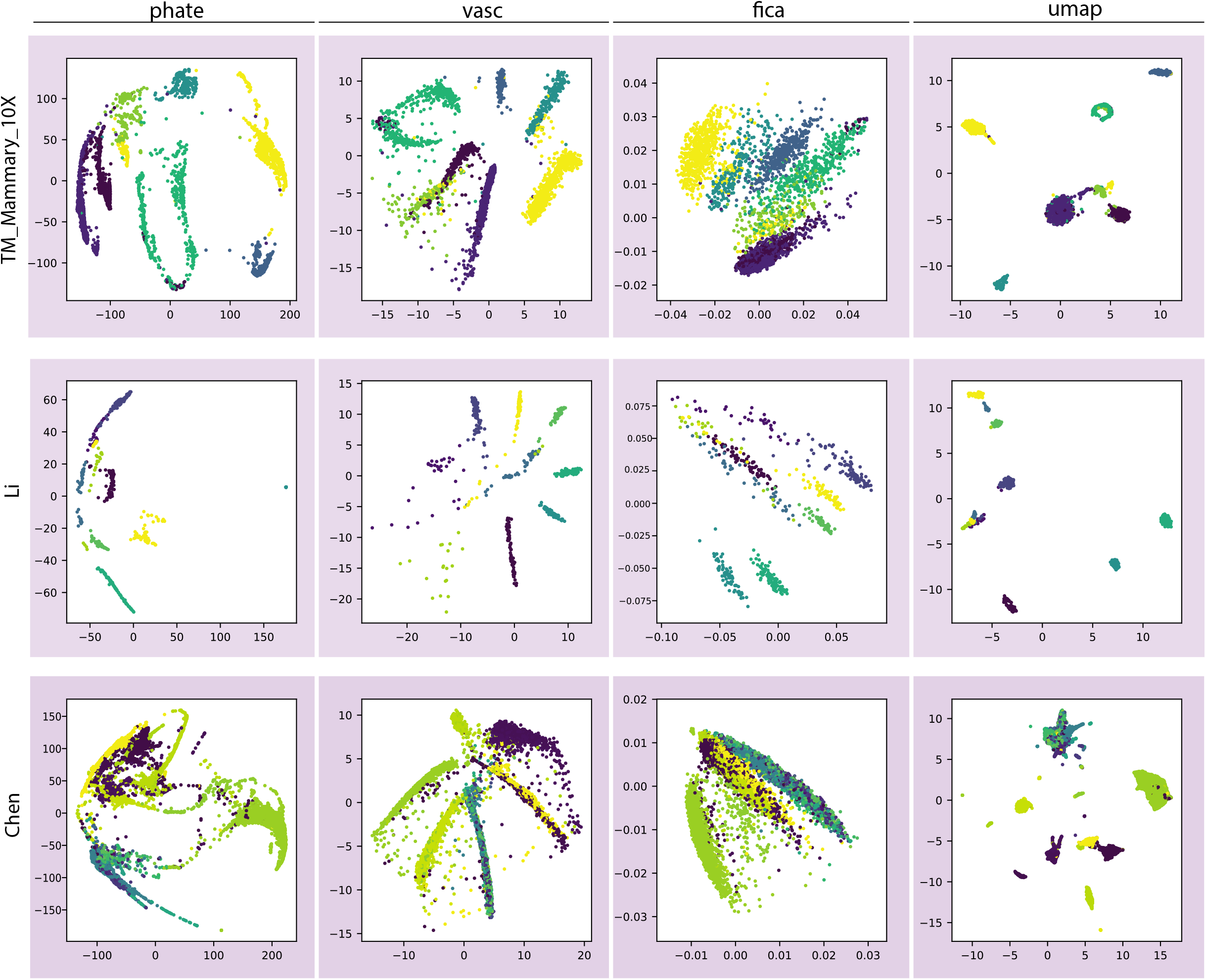
Visualization of selected dimensionality reduction methods across a few datasets which obtained high ARI during DBSCAN analysis (*c.f*. **Fig. 3**).

### Memory requirements and compute time

Given the ever-increasing sample sizes of scRNA-seq data, an important question is how computational requirements scale. Thus, runtime and memory usage were recorded for each low dimension embedding for 31 of our methods, in our real datasets ranging from 66-27,499 cells (Supplementary Fig. 7-8).

Despite the relative variation in resource requirements between methods, time and memory requirements grew approximately linearly or log-linearly with respect to sample size. Slower growth trends (*i.e*. log-linear) were observed in the compute time for methods which relied heavily on random sampling of the data as opposed to full-matrix calculations, *e.g*., VPAC, UMAP, and PCA (which utilized a random-projection algorithm in our sklearn implementation).

How the number of dimensions affected computational resources was method specific. While runtime increased approximately proportionally to dimension number for most methods (*e.g*. BD and PMF), others were minimally affected (*e.g*. most PCA-like methods, Supplementary Fig. 7). In contrast, memory usage generally did not scale with the number of embeddings, except with ZIFA and PSMF (Supplementary Fig. 8).

Notably, PHATE and LDA had average computational requirements. In particular, PHATE employs a sub-sampling approach which allows it to scale well to large datasets. Thus, their greater performance in clustering and IVM analyses is not compromised by intractable runtime or memory usage, making them attractive options for dimensionality reduction.

## Discussion

Here, we have performed a comprehensive benchmarking of dimensionality reduction methods for scRNA-seq data. We examined 33 methods, many of which were yet to be independently assessed, and a summary of their performance is provided in Fig. 6. Notably, two of the best performing methods – LDA and PHATE – were not assessed in previous benchmarks^13–15^, highlighting the importance of our broad-ranging benchmarking strategy, as well as the growth in the number of methods in recent years.

**Fig. 6:**
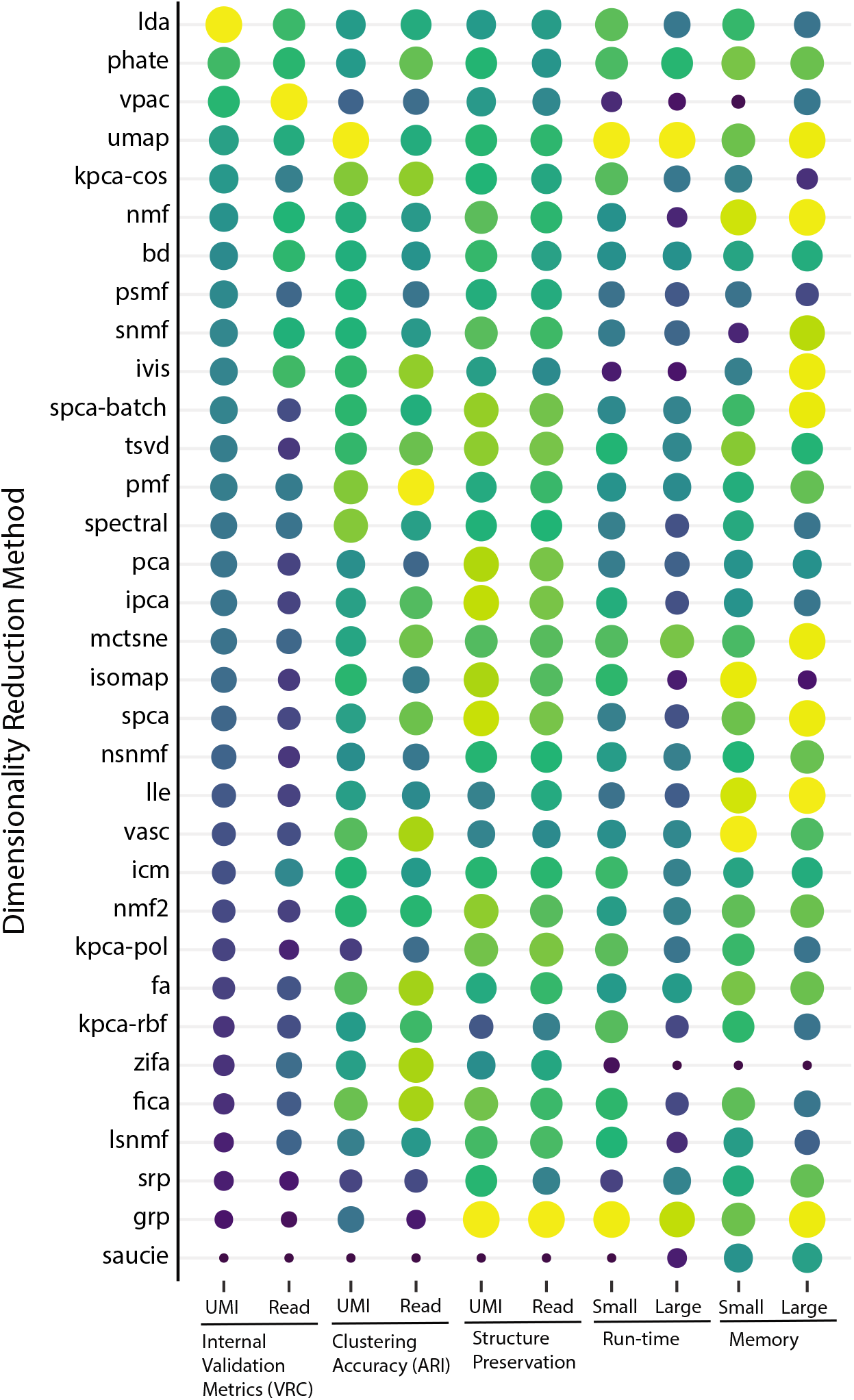
Performance summary of dimensionality reduction methods. (row) according to the assessed criteria. Larger dots indicate greater relative performance for a given metric (*i.e*., column). From left to right: ranking of VRC on UMI/read datasets, ARI on UMI/read datasets, preservation of global structure on UMI/read datasets, and time/memory requirements for an exemplar small (Darmanis: 466 cells) and large (Shekar: 27,499 cells) dataset. The absence of a dot indicates that data was not recorded.

We employed an approach to benchmarking of dimensionality reduction methods which relies solely on the use of IVMs to quantify how well formed the existing clusters appear in the low-dimensional embedding. The primary benefit of this approach is that it is not reliant on downstream analyses which may confound benchmarking results. To the best of the authors’ knowledge, it is the only such method which may also take advantage of gold-standard, or labelled data.

Our results suggest that several overlooked dimensionality reduction methods may yield better performance than the current popular methods like UMAP or t-SNE, or those recommended in past benchmarking studies such PCA or FA^13^. We found that bypassing downstream analyses may provide better power to differentiate the performance of dimensionality reduction methods. Latent Dirichlet Allocation (LDA) was found to consistently produce low-dimension embeddings with the best internal validation measures. When considering the use of IVMs in unsupervised DBSCAN clustering, PHATE was found to result in the highest Adjusted Rand Index (with neither of these methods being benchmarked by Sun *et al*. (2019)). Several other secondary measures of performance were also considered. We found that Gaussian Random Projection (GRP) and each of the various principal component methods did the best job at preserving the global structure of the data. This is likely due to the highly linear nature of such methods. While Heiser & Lau (2020) found similar results, this does contrast to past benchmarking work by Sun *et al*. (2019), which found PCA had average performance; the discrepancy may result from the use of different metrics, with Sun *et al*. (2019) using a Jaccard Index on sets of nearest neighbors, rather than a global correlation.

In terms of visualization, methods which performed well in DBSCAN clustering also tended to provide embeddings in two dimensions which clearly distinguish between cell-types, despite the unsupervised approach.

Based on our results there are a number of recommendations we can provide to researchers and method developers. Internal validation measures, particularly Variance Ratio Criterion (VRC), should be used in the presence of labelled data to guide the development of dimensionality reduction methods without the need for potentially computationally intensive downstream analyses. Furthermore, IVMs can successfully be used as an unsupervised method to predict an optimal clustering. Our results suggest that it may be worth researchers considering alternate dimensionality reduction methods depending on the desired result. Methods such as LDA or PHATE, despite their relative obscurity in scRNA-seq analysis pipelines, are promising alternatives to traditionally used or recommended methods; however, if preservation of global structure is a priority, linear methods such as GRP or PCA are better candidates.

Where possible, we followed the guidelines of Weber *et al*. regarding best practice benchmarking procedures^18^. This includes, but is not limited to: a survey of existing and relevant methods, using default parameters for all packages in order to avoid implicit bias, selection of a wide range of representative scRNA-seq datasets, evaluation of performance according to several key quantitative metrics, evaluation of secondary measures, and the documentation and publishing of all relevant data and code to enable reproducibility.

This study has some limitations. Many of the datasets used did not possess “gold-standard” labels; instead, labels predicted by the publishing group were used to assess performance. However, we found that most methods’ performance did not differ across the different approaches for determining cell-type labels, helping to address this concern (Supplementary Fig. 3). Furthermore, even without correcting for multiple comparisons there were few significant differences in methods nearby in rank (Fig. 2b and Fig. 4b); these rankings should therefore be considered as a general guide and not absolute. Finally, while generally consistent, the rankings of methods in both the IVM and clustering analyses are not in complete agreement, which may be due to the poor resolving-power of the clustering analysis. Hence, it is still important to benchmark methods for specific downstream analyses and usecases.

## Methods

### Dimensionality reduction method selection

To determine which methods should be included in this benchmark, a survey was conducted on the software packages listed at scRNA-tools.org. Only packages listed under the dimensionality reduction category were considered. The documentation and source code of each package was inspected to determine which dimensionality reduction methods were utilized. A summary of this survey is available in Supplementary Table 4. Only dimensionality reduction methods with open-source implementations available in Python 3 were selected for inclusion in this benchmark. Methods were excluded if they did not possess sufficient documentation, raised difficult to fix errors at runtime, or only exposed a command-line interface. The final list of 33 dimensionality reduction methods is summarized in Table 1.

### Datasets

55 datasets were obtained from a curated set of publicly available scRNA-seq datasets maintained by the Hemberg Lab^24^. Each of these datasets contained cell-type assignments which were used as the ‘ground-truth’ for measures of accuracy and internal validation. This is a diverse collection of datasets which spans a wide range of sequencing protocols, tissue types, and number of cells. The properties of each dataset are summarized in Supplementary Table 1.

### Generation of low dimension embeddings

Each method was applied to each dataset to generate embeddings in 2, 4, 8, 16, 32, 48, and 96 dimensions for both log1p-transformed and non-transformed data after count-per-million normalization. In accordance with the benchmark guidelines set out by Weber *et al*. (2019), no method was given special treatment^18^. Default arguments were used for all methods where applicable.

### Internal Validation Measures (IVMs)

IVMs were used to assess how well-defined the clusters appear after dimensionality reduction. Three measures implemented by sci-kit learn (version 0.22.1) were used for this purpose: silhouette score (SS), variance ratio criterion (VRC), and Davies Bouldin index (DBI)^25^. Additionally, SS was separately calculated using Euclidean, standardized Euclidean, correlation, and cosine similarity measures. The cell-type predictions supplied with each dataset were used to estimate the true clusters.

#### Silhouette Score

SS is calculated as the mean silhouette coefficient over the dataset, and varies between −1 and 1 with larger values being better^20^. A high silhouette score indicates that each point is more similar to points in its own cluster than to points from other clusters. Assume that data have been clustered via any technique into *k* clusters. For each point *i* in cluster *C_i_* (i.e., *i* ∈ *C_i_* assuming |*C_i_*| > 1), the silhouette coefficient is defined as:

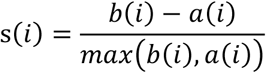

Where

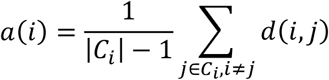

is the average distance, *d*(*i, j*), of point *i* to each other point within the same cluster, C*_i_*, and

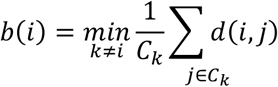

is the average nearest-neighbor distance to each cluster.

#### Variance Ratio Criterion

VRC, also known as Calinski Harabasz score, is the ratio of between-cluster dispersion to within-cluster dispersion and is defined as^21^:

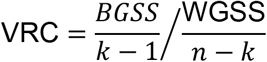

Where *k* is the number of clusters, *n* is the number of data points, BGSS is the between group sum-of-squares, and WGSS is the within group sum-of-squares. Larger values of VRC indicate high dispersion between clusters and low dispersion within clusters.

#### Davies Bouldin Index

DBI is the average similarity of each cluster with its closest neighbor cluster^22^. It is defined as:

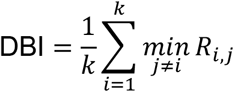

Where *k* is the number of clusters, and *R_i,j_* is the similarity of clusters *i* and *j* which is defined as:

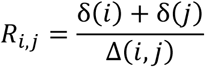

With δ(*i*) being the average distance of each point in cluster *i* to its corresponding centroid, and Δ(*i, j*) being the distance between the two centroids of the two clusters.

### Statistical analysis of Internal Validation Measures

#### Intra-IVM and intra-dataset comparisons

Rank-based analyses were performed in order to make minimal assumptions about the distribution of IVMs as well as to allow for comparison between SS, DBI, and VRC. Dimensionality reduction methods were ranked on each dataset according to each of the three IVMs, and median rank across datasets were computed for each method. These median ranks were again ranked to compute concordance between IVMs.

Kendall’s W was calculated using the Synchrony^26^ package to measure the concordance in the ranking of dimensionality reduction methods by each of the various internal validation measures. Kendall’s W is calculated as:

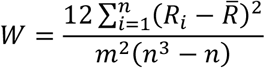

Where *R*_i_ is the sum of ranks for the i^th^ method, 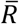 is the average *R* across all methods, *n* is the number of variables (methods), and *m* is the number of judges (IVMs: VRC, SS, DBS). Concordance was also calculated between datasets, and similarity measures (Euclidean, standardized Euclidean, cosine, correlation). Permutation testing was used to estimate p-values.

#### Dimensionality reduction method comparisons

Pairwise Sign tests were used to test for significant differences between methods for each IVM. The sign test is a non-parametric test of whether one distribution is stochastically greater than the other. It relies on the fact that under the null hypothesis of no difference, there is an equal probability of observing a better result from either method. Therefore, the number of times one method outperforms another is binomially distributed, thus allowing for a direct calculation of a p-value. P-values were not corrected for multiple comparisons in order to maximize the power of this test to detect possible differences between methods.

### Clustering by DBSCAN random search

#### DBSCAN optimization

Density-based clustering was performed over the four distance measures for each method by a random search over the hyper-parameter space of the DBSCAN algorithm implemented in sci-kit learn^17,25^. DBSCAN requires two parameters: epsilon (*ϵ*), which determines the radius within which other points are considered “neighbors”; and minimum points (minPts), which is the minimum number of points within a cluster. We opted for random search as opposed to grid search for its theoretical guarantees regarding convergence^27^. We used Monte Carlo random sampling to generate 2,000 random trials for each embedding and distance measure. First, minPts is randomly sampled such that minPts is uniformly distributed along a logarithmic scale with a minimum of 5 and maximum of 15% the size of the dataset.

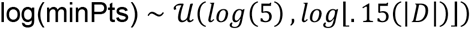

Where 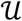 represents the uniform distribution and |*D*| is the size of the dataset. The logarithms ensure a more balanced exploration of the sample space because, for example, changing minPts from 5 to 6 will generally have a larger effect on clustering than changing from 50 to 51. Next, we sampled ϵ according to a truncated exponential distribution:

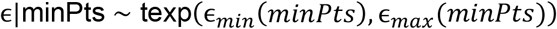

Where *ϵ_min_*(*minPts*) is determined by the minimum radius required for at least one point in the dataset to have minPts neighbors, and *ϵ_max_*(*minPts*) is the distances at which all points have minPts neighbours. This distribution is chosen in order to maximize the exploration of areas which are likely to be sensitive to changes in the parameter. That is, when ϵ is large, small changes will have no effect on the clustering, however, when ϵ is small, there will be a much larger effect on the final clustering.

For each random trial, we calculated IVMs for the resulting clustering. These IVMs were used to determine the optimal hyper-parameters and dimension of embedding for each dimensionality reduction method. To assess the accuracy of clustering, we calculated the adjusted rand index (ARI), and normalized mutual information (NMI) comparing true cell labels and inferred cell labels obtained by DBSCAN.

#### Comparing optimization approaches

DBSCAN optimization was performed using Euclidean, standardized Euclidean, correlation, and cosine distance measures in addition to the three IVMs. Therefore, there are 12 possible combinations of IVM and distance measure which could be used for clustering. These were compared by considering the average ARI of each method across all datasets. The combination of VRC and standardized Euclidean distance was observed to yield the maximum ARI for most methods (14 out of 33). To test that this was not due to random chance, the probability of observing this extreme of an observation was estimated over 100,000 Monte Carlo simulations of a multivariate hypergeometric distribution (p < 0.001).

#### Dimensionality reduction method comparisons

Both pairwise Wilcoxon Sign-Rank and pairwise Sign tests were employed to compare the performance of each method across the 55 datasets. P-values were not corrected for multiple comparisons in order to maximize the power of this test to detect differences between methods.

#### Comparison with IVM analysis

The median ranks across datasets for each method from the IVM analysis were compared with median ranks from this clustering analysis. Both Pearson’s correlation and Kendall’s W were calculated to compare the two rankings.

### Preservation of global structure analysis

For each full-dataset and low-dimension embedding, pairwise distances were calculated for the following similarity measures: Euclidean, standardized Euclidean, correlation, and cosine. To test whether global structure was well preserved, Spearman correlations were calculated between the full-dataset and low-dimension embedding, the idea being that the relative distance between points should not change after dimensionality reduction if global structure is well preserved.

### Memory requirements and compute time

Generation of embeddings for all methods except for ZIFA and GPU based methods were run on 4 cores of a 2.8 Ghz X5660 with up to 64GB of RAM. ZIFA embeddings were run on 8 cores of a 2.2 Ghz E5-4620 with up to 128GB of RAM. GPU methods (saucie and VASC) were run on an Nvidia Quadro M5000. During computation of low-dimension embeddings, total runtime (s) and maximum resident set size (kb) were recorded.

## Code availability

Code for this study is available from github.com/ForrestCKoch/scRNA-Dimensionality-Reduction and is documentation is documented at scrna-dimensionality-reduction.rtfd.io/.

## Notes

https://github.com/ForrestCKoch/scRNA-Dimensionality-Reduction

### Competing Interest Statement

The authors have declared no competing interest.

### Summary of Updates

- Author names and institutions were revised - License changed to CC-NC-ND

