## Supplemental Tables 1-3 for "Supervised Application of Internal Validation Measures to Benchmark Dimensionality Reduction Methods in scRNA-seq Data"

### Supplementary Tables

***Supplementary Table 1** Summary of datasets used in benchmarking.*

| Datasets | Protocol | Accession | Species | Tissue type | Cells | Genes | Cell types | Min cell type% | Max cell type% | UMI/ read | Cell type determination |
|---|---|---|---|---|---|---|---|---|---|---|---|
| **baron human[1]** | inDrop | GSE84133 | human | pancreas | 8569 | 20125 | 14 | 0.08 | 0.21 | UMI | Expression-based |
| **baron mouse[1]** | inDrop | GSE84133 | mouse | pancreas | 1886 | 14878 | 13 | 0.32 | 11.56 | UMI | Expression-based |
| **campbell[2]** | Drop-seq | GSE93374 | mouse | brain | 21086 | 26774 | 35 | 0.23 | 2.50 | UMI | Dimension-reduction-based |
| **chen[3]** | Drop-seq | GSE87544 | mouse | brain | 14437 | 23284 | 47 | 0.11 | 0.26 | UMI | Dimension-reduction-based |
| **darmanis[4]** | SMARTer | GSE67835 | human | brain | 466 | 22088 | 9 | 3.43 | 8.15 | Read | Dimension-reduction-based |
| **deng reads[5]** | Smart-Seq2 | GSE45719 | mouse | embryo | 268 | 22431 | 6 | 4.48 | 13.81 | Read | Gold |
| **deng rpkms[5]** | Smart-Seq2 | GSE45719 | mouse | embryo | 268 | 22431 | 6 | 4.48 | 13.81 | Read | Gold |
| **fan[6]** | SUPeR-seq | GSE53386 | mouse | embryo | 66 | 26357 | 6 | 10.61 | 33.33 | Read | Gold |
| **goolam[7]** | Smart-Seq2 | E-MTAB-3321 | mouse | embryo | 124 | 41480 | 5 | 4.84 | 25.81 | Read | Gold |
| **klein[8]** | inDrop | GSE65525 | mouse | embryo stem cells | 2717 | 24175 | 4 | 11.15 | 34.34 | UMI | Gold |
| **lake[9]** | C1 | phs000833.v3.p1 | human | brain | 3042 | 25051 | 16 | 1.48 | 2.30 | read | Expression-based |
| **li[10]** | SMARTer | GSE81861 | human | colorectal tumor | 561 | 55186 | 9 | 4.10 | 12.30 | Read | Gold |
| **manno human[11]** | STRT-Seq UMI | GSE76381 | human | brain | 4029 | 20560 | 56 | 0.12 | 0.45 | UMI | Expression-based |
| **manno mouse[11]** | STRT-Seq UMI | GSE76381 | mouse | brain | 2150 | 24378 | 32 | 0.56 | 1.26 | UMI | Expression-based |
| **marques[12]** | C1 | GSE75330 | mouse | brain | 5053 | 23556 | 13 | 1.50 | 9.12 | Read | Expression-based |
| **muraro[13]** | CEL-Seq2 | GSE85241 | human | pancreas | 2126 | 19127 | 10 | 0.14 | 10.30 | YNU | Expression-based |
| **pollen[14]** | SMARTer | SRP041736 | human | neonatal-foreskin | 301 | 23730 | 11 | 2.33 | 17.94 | Read | Gold |
| **romanov[15]** | C1 | GSE74672 | mouse | brain | 2881 | 24341 | 7 | 1.67 | 2.46 | Read | Expression-based |
| **segerstolpe[16]** | Smart-Seq2 | E-MTAB-5061 | human | pancreas | 3514 | 25525 | 15 | 0.23 | 0.72 | Read | Dimension-reduction-based |
| **shekhar[17]** | Drop-seq | GSE81904 | mouse | retina | 27499 | 13166 | 6 | 0.18 | 0.94 | UMI | Dimension-reduction-based |
| **Bladder 10X[18]** | Chromium | GSE132042 | mouse | bladder | 2500 | 23433 | 5 | 2.28 | 10.64 | Umi | Dimension-reduction-based |
| **Bladder FACS[18]** | Smart-Seq2 | GSE132042 | mouse | bladder | 1638 | 23433 | 4 | 6.04 | 40.05 | Read | Dimension-reduction-based |
| **Brain Microglia FACS[18]** | Smart-Seq2 | GSE132042 | mouse | microglia | 4762 | 23433 | 3 | 0.76 | 90.91 | Read | Dimension-reduction-based |
| **Brain Neurons FACS[18]** | Smart-Seq2 | GSE132042 | mouse | neurons | 5799 | 23433 | 11 | 0.25 | 3.59 | Read | Dimension-reduction-based |
| **Colon FACS[18]** | Smart-Seq2 | GSE132042 | mouse | colon | 4149 | 23433 | 6 | 0.60 | 19.62 | Read | Dimension-reduction-based |
| **Fat FACS[18]** | Smart-Seq2 | GSE132042 | mouse | fat | 5862 | 23433 | 11 | 0.46 | 3.77 | Read | Dimension-reduction-based |
| **Heart 10X[18]** | Chromium | GSE132042 | mouse | heart | 654 | 23433 | 7 | 3.21 | 33.94 | UMI | Dimension-reduction-based |
| **Heart FACS[18]** | Smart-Seq2 | GSE132042 | mouse | heart | 7115 | 23433 | 11 | 0.16 | 17.95 | Read | Dimension-reduction-based |
| **Kidney 10X[18]** | Chromium | GSE132042 | mouse | kidney | 2782 | 23433 | 9 | 1.62 | 4.06 | UMI | Dimension-reduction-based |
| **Kidney FACS[18]** | Smart-Seq2 | GSE132042 | mouse | kidney | 865 | 23433 | 7 | 4.39 | 7.51 | Read | Dimension-reduction-based |
| **Liver 10X[18]** | Chromium | GSE132042 | mouse | liver | 1924 | 23433 | 3 | 1.04 | 52.29 | UMI | Dimension-reduction-based |
| **Liver FACS[18]** | Smart-Seq2 | GSE132042 | mouse | liver | 981 | 23433 | 6 | 2.96 | 19.98 | Read | Dimension-reduction-based |
| **Lung FACS[18]** | Smart-Seq2 | GSE132042 | mouse | lung | 1923 | 23433 | 17 | 0.10 | 0.68 | Read | Dimension-reduction-based |
| **Mammary 10X[18]** | Chromium | GSE132042 | mouse | mammary | 4481 | 23433 | 7 | 4.15 | 17.05 | UMI | Dimension-reduction-based |
| **Mammary FACS[18]** | Smart-Seq2 | GSE132042 | mouse | mammary | 2663 | 23433 | 5 | 1.84 | 16.07 | Read | Dimension-reduction-based |
| **Marrow 10X[18]** | Chromium | GSE132042 | mouse | marrow | 4112 | 23433 | 9 | 1.53 | 3.94 | UMI | Dimension-reduction-based |
| **Marrow FACS[18]** | Smart-Seq2 | GSE132042 | mouse | marrow | 5355 | 23433 | 9 | 2.54 | 34.51 | Read | Dimension-reduction-based |
| **Muscle 10X[18]** | Chromium | GSE132042 | mouse | muscle | 4543 | 23433 | 9 | 7.28 | 26.26 | UMI | Dimension-reduction-based |
| **Muscle FACS[18]** | Smart-Seq2 | GSE132042 | mouse | muscle | 2102 | 23433 | 8 | 1.67 | 21.03 | Read | Dimension-reduction-based |
| **Pancreas FACS[18]** | Smart-Seq2 | GSE132042 | mouse | pancreas | 1961 | 23433 | 10 | 1.48 | 20.96 | Read | Dimension-reduction-based |
| **Skin FACS[18]** | Smart-Seq2 | GSE132042 | mouse | skin | 2464 | 23433 | 5 | 1.83 | 55.64 | Read | Dimension-reduction-based |
| **Spleen 10X[18]** | Chromium | GSE132042 | mouse | spleen | 9573 | 23433 | 5 | 1.50 | 23.89 | UMI | Dimension-reduction-based |
| **Spleen FACS[18]** | Smart-Seq2 | GSE132042 | mouse | spleen | 1718 | 23433 | 4 | 2.79 | 23.46 | Read | Dimension-reduction-based |
| **Thymus 10X[18]** | Chromium | GSE132042 | mouse | thymus | 1431 | 23433 | 3 | 7.83 | 92.03 | UMI | Dimension-reduction-based |
| **Thymus FACS[18]** | Smart-Seq2 | GSE132042 | mouse | thymus | 1580 | 23433 | 3 | 2.09 | 79.11 | Read | Dimension-reduction-based |
| **Tongue 10X[18]** | Chromium | GSE132042 | mouse | tongue | 7538 | 23433 | 3 | 32.21 | 67.79 | UMI | Dimension-reduction-based |
| **Tongue FACS[18]** | Smart-Seq2 | GSE132042 | mouse | tongue | 1432 | 23433 | 3 | 25.21 | 72.14 | Read | Dimension-reduction-based |
| **Trachea FACS[18]** | Smart-Seq2 | GSE132042 | mouse | trachea | 1391 | 23433 | 5 | 2.37 | 40.91 | Read | Dimension-reduction-based |
| **tasic reads[19]** | SMARTer | GSE71585 | mouse | brain | 1679 | 24150 | 18 | 0.54 | 16.38 | Read | Dimension-reduction-based |
| **tasic rpkms[19]** | SMARTer | GSE71585 | mouse | brain | 1679 | 24057 | 18 | 0.54 | 1.43 | Read | Dimension-reduction-based |
| **treutlein[20]** | SMARTer | GSE52583 | mouse | lung | 80 | 23271 | 5 | 3.75 | 15.00 | Read | Expression-based |
| **usoskin[21]** | STRT-Seq | GSE59739 | mouse | brain | 622 | 25334 | 4 | 13.02 | 37.46 | Read | Dimension-reduction-based |
| **xin[22]** | SMARTer | GSE81608 | human | pancreas | 1600 | 39851 | 8 | 3.28 | 59.38 | Read | Expression-based |
| **yan[23]** | Tang | GSE36552 | human | embryo | 90 | 20214 | 6 | 6.67 | 17.78 | Read | Gold |
| **zeisel[24]** | STRT-seq | GSE60361 | mouse | brain | 3005 | 19972 | 9 | 0.87 | 6.59 | UMI | Expression-based |

***Supplementary Table 2***. *Dimensionality reduction methods were ranked according to each internal validation measure (IVM). This table shows median rank across all datasets for each IVM. DBS – Davies Bouldin Score, SS_(COR/COS/EUC/SEU) silhouette score (correlation/cosine/Euclidean/standardized Euclidean) distances, VRC – variance ratio criterion. Rows are sorted by their average median rank across DBS, SS_EUC, and VRC (2nd far right column). The far-right column shows the average across all IVMs (including SS variations).*

| measure | DBI | SS-COR | SS-COS | SS-EUC | SS-SEU | VRC | Overall | Overall* |
|---|---|---|---|---|---|---|---|---|
| **lda** | 7 | 4 | 3 | 3 | 6 | 2 | 4.00 | 4.17 |
| **vpac** | 9 | 10 | 10 | 2 | 4 | 5 | 5.33 | 6.67 |
| **phate** | 7 | 10 | 10 | 5 | 6 | 6 | 6.00 | 7.33 |
| **nmf-nnsvd** | 8 | 5 | 5 | 7 | 7 | 6 | 7.00 | 6.33 |
| **ivis** | 6 | 12 | 10 | 9 | 9 | 8 | 7.67 | 9.00 |
| **icm** | 9 | 7 | 6 | 8 | 9 | 7 | 8.00 | 7.67 |
| **snmf** | 13 | 9 | 7 | 10 | 9 | 7 | 10.00 | 9.17 |
| **kpca-cos** | 8 | 11 | 11 | 9 | 11 | 13 | 10.00 | 10.50 |
| **fa** | 11 | 11 | 10 | 8 | 10 | 14 | 11.00 | 10.67 |
| **zifa** | 13 | 14 | 13 | 12 | 13 | 14 | 13.00 | 13.17 |
| **psmf** | 14 | 8 | 8 | 13 | 17 | 14 | 13.67 | 12.33 |
| **vasc** | 11 | 20 | 19 | 13 | 13 | 17 | 13.67 | 15.50 |
| **fica** | 12 | 18 | 16 | 12 | 13 | 18 | 14.00 | 14.83 |
| **spca** | 11 | 12 | 14 | 13 | 12 | 19 | 14.33 | 13.50 |
| **bd** | 21 | 17 | 18 | 24 | 9 | 7 | 15.00 | 16.00 |
| **pca** | 13 | 18 | 19 | 16 | 13 | 20 | 16.33 | 16.50 |
| **ipca** | 14 | 18.5 | 19 | 16 | 13 | 19.5 | 16.50 | 16.67 |
| **lsnmf** | 19 | 16 | 16 | 18 | 21 | 14 | 17.00 | 17.33 |
| **tsvd** | 14 | 9 | 7 | 17 | 15 | 21 | 17.33 | 13.83 |
| **pmf** | 17 | 18 | 14 | 15 | 19 | 13 | 17.33 | 16.00 |
| **tsne** | 16 | 24 | 25 | 19 | 21 | 17 | 17.33 | 20.33 |
| **nsnmf** | 22 | 17 | 16 | 19 | 21 | 18 | 17.67 | 18.83 |
| **spca-batch** | 17 | 19 | 19 | 19 | 19 | 19 | 18.33 | 18.67 |
| **spectral** | 22 | 25 | 23 | 19 | 21 | 14 | 18.33 | 20.67 |
| **umap** | 24 | 21 | 22 | 18 | 17 | 11 | 19.67 | 18.83 |
| **kpca-rbf** | 16 | 17 | 20 | 22 | 21 | 24 | 20.67 | 20.00 |
| **lle** | 26 | 21 | 20 | 20 | 21 | 19 | 21.67 | 21.17 |
| **isomap** | 24 | 25 | 26 | 24 | 24 | 22 | 23.33 | 24.17 |
| **nmf-lee** | 27 | 23 | 21 | 25 | 26 | 20 | 24.00 | 23.67 |
| **kpca-pol** | 21 | 23 | 25 | 26 | 22 | 27 | 24.67 | 24.00 |
| **srp** | 29 | 25 | 26 | 29 | 30 | 29 | 29.00 | 28.00 |
| **grp** | 29 | 29 | 29 | 29 | 29 | 30 | 29.33 | 29.17 |
| **saucie** | 32 | 32 | 32 | 32 | 32 | 32 | 32 | 32.00 |

***Supplementary Table 3***. *Average ARI across datasets for DRM/DBSCAN optimization method. * - max ARI in column (method). †- p < 0.05 for sign-test against VRC-SEU, § - p < 0.05 for paired Wilcoxon test against VRC.SEU*

| measure | phate | vasc | fica | umap | zifa | lsnmf | kpca-cos | nmf-lee | kpca-sig | pca | spca | fa | tsne | ipca | snmf | tsvd | spectral | pmf | ivis | icm | lda | kpca-rbf | bd | lle | isomap | spca-batch | nmf-lee | nsnmf | vpac | kpca-pol | grp | srp | saucie | overall |
|---|---|---|---|---|---|---|---|---|---|---|---|---|---|---|---|---|---|---|---|---|---|---|---|---|---|---|---|---|---|---|---|---|---|---|
| **VRC-EUC** | .53 * | .46 | .45 *† | .41 | .42 | .41 | .40 | .40 * | .30 | .37 | .38 | .39 * | .38 | .36 | .36 | .34 | .34 | .33 | .32 | .32 | .34 | .30 | .25 | .28 | .29 | .31 | .28 * | .28 * | .27 *† | .20 | .15 | .10 | .00 | .32 |
| **VRC-SEU** | .53 | .46 * | .45 | .44 * | .43 * | .42 * | .40 | .40 | .30 | .39 * | .39 * | .36 | .39 * | .38 * | .37 * | .36 * | .34 | .34 | .35 * | .32 | .34 * | .31 | .31 | .28 | .30 | .31 * | .26 | .27 | .27 | .22 * | .16 | .09 | .00 | .33 |
| **VRC-COS** | .50 | .46 | .39 | .42 | .39 | .36 | .41 * | .36 | .36 | .32 | .32 | .38 | .38 | .31 | .35 | .28 | .34 | .30 | .34 | .30 | .33 | .33 * | .33 * | .29 | .32 * | .30 | .24 | .26 | .27 | .21 | .16 * | .16 *†§ | .00 | .32 |
| **VRC-COR** | .50 | .27 | .37 | .41 | .34 | .25 | .39 | .31 | .33 | .28 | .28 | .39 | .36 | .25 | .29 | .30 | .30 | .24 | .33 | .34 * | .30 | .34 | .28 | .32 *† | .28 | .28 | .21 | .25 | .27 | .20 | .15 | .15 | .00 | .29 |
| **SS-EUC** | .43 | .41 | .16 | .38 | .36 | .38 | .39 | .17 | .20 | .22 | .21 | .17 | .22 | .21 | .12 | .17 | .34 | .24 | .25 | .06 | .34 | .06 | .12 | .23 | .18 | .12 | .13 | .27 | .26 | .03 | .07 | .06 | .00 | .21 |
| **SS-SEU** | .26 | .42 | .16 | .36 | .31 | .38 | .33 | .20 | .18 | .15 | .16 | .15 | .27 | .17 | .22 | .15 | .34 | .28 | .26 | .08 | .30 | .06 | .19 | .25 | .16 | .15 | .14 | .23 | .27 | .06 | .06 | .02 | .00 | .20 |
| **SS-COS** | .51 | .44 | .35 | .37 | .39 | .35 | .40 | .30 | .40 *†§ | .32 | .33 | .36 | .09 | .31 | .29 | .26 | .35 *† | .35 *† | .33 | .28 | .32 | .32 | .08 | .21 | .31 | .29 | .25 | .27 | .26 | .12 | .10 | .12 | .00 | .29 |
| **SS-COR** | .20 | .22 | .17 | .22 | .23 | .03 | .21 | .15 | .21 | .18 | .15 | .21 | .18 | .16 | .16 | .19 | .16 | .21 | .22 | .30 | .24 | .05 | .29 | .17 | .18 | .18 | .14 | .18 | .27 | .09 | .05 | .10 | .00 | .17 |
| **DBI-EUC** | .19 | .17 | .15 | .34 | .25 | .16 | .23 | .17 | .15 | .18 | .18 | .17 | .29 | .15 | .15 | .16 | .26 | .18 | .22 | .10 | .30 | .12 | .11 | .19 | .14 | .15 | .15 | .15 | .27 | .11 | .08 | .05 | .00 | .17 |
| **DBI-SEU** | .18 | .19 | .15 | .33 | .25 | .18 | .22 | .18 | .16 | .17 | .17 | .16 | .25 | .15 | .17 | .16 | .27 | .18 | .22 | .13 | .31 | .12 | .15 | .20 | .12 | .15 | .17 | .18 | .27 | .15 | .08 | .04 | .00 | .18 |
| **DBI-COS** | .23 | .28 | .18 | .33 | .26 | .24 | .27 | .20 | .22 | .17 | .16 | .20 | .29 | .16 | .20 | .18 | .28 | .18 | .23 | .19 | .30 | .12 | .21 | .21 | .17 | .11 | .14 | .21 | .27 | .15 | .07 | .08 | .00 | .20 |
| **DBI-COR** | .21 | .16 | .13 | .34 | .20 | .28 | .22 | .22 | .20 | .15 | .11 | .13 | .30 | .14 | .22 | .14 | .16 | .18 | .17 | .24 | .30 | .14 | .19 | .19 | .17 | .07 | .15 | .2 | .27 | .17 | .08 | .06 | .00 | .18 |
| **MAX** | .53 | .46 | .45 | .44 | .43 | .42 | .41 | .4 | .4 | .39 | .39 | .39 | .39 | .38 | .37 | .36 | .35 | .35 | .35 | .34 | .34 | .34 | .33 | .32 | .32 | .31 | .28 | .28 | .27 | .22 | .16 | .16 | .00 | .33 |
| **RANK** | 1 | 2 | 3 | 4 | 5 | 6 | 7 | 8 | 9 | 10 | 11 | 12 | 13 | 14 | 15 | 16 | 17 | 18 | 19 | 20 | 21 | 22 | 23 | 24 | 25 | 26 | 27 | 28 | 29 | 30 | 31 | 32 | 33 | – |
