## Supplemental Table 4 for "Supervised Application of Internal Validation Measures to Benchmark Dimensionality Reduction Methods in scRNA-seq Data"

*Supplementary Table 4. The documentation and source code of scRNA-tools.org packages as summarized below were inspected to determine which dimensionality reduction methods were utilized.*

| Package | Dimensionality Reduction | DOI/Reference Link | GitHub Repo/Link |
|---|---|---|---|
| Altanalyze | pca,svd,t-sne | 10.1038/nature19348 | nsalomonis/altanalyze |
| BigSCale2 | umap,tsne | 10.1186/s13059-019-1713-4 | iaconogi/bigSCale2 |
| ccfindr | nmf | 10.26508/lsa.201900443 | hjunwoo/ccfindR |
| celda | lda | N/A | campbio/celda |
| CellRouter | pca,tsne for visualization | 10.1038/s41467-018-03214-y | edroaldo/cellrouter |
| Cell Ranger | N/A | N/A | 10XGenomics/cellranger |
| Cell Trails | spectral,tsne,pca | 10.1016/j.celrep.2018.05.002 | dcellwanger/CellTrails |
| CIDR | pca | N/A | VCCRI/CIDR |
| cNMF | nmf | 10.7554/eLife.43803 | dylkot/cNMF |
| conos | common pca, joint nmf | 10.1038/s41592-019-0466-z | hms-dbmi/conos |
| csmf | nmf | 10.1101/272443 | page.amss.ac.cn/shihua.zhang/software.html |
| cyclum | autoencoder | 10.1101/625566 | KChen-lab/cyclum |
| dca_r | dca,pca | 10.1371/journal.pcbi.1006391 | cran/DCA |
| destiny path | pca,elastic embedding | 10.1093/bioinformatics/bty1009 | ucasdp/DensityPath |
| destiny | diffusion maps | N/A | theislab/destiny |
| dhaka | variational autoencoder | 10.1101/183863 | MicrosoftGenomics/Dhaka |
| DoubletFinder | pca | 10.1016/j.cels.2019.03.003 | chris-mcginnis-ucsf/DoubletFinder |
| DropClust | pca,umap | 10.1101/596924 | debsin/dropClust |
| DrSeq2 | tsne,pca,simlr | 10.1371/journal.pone.0180583 | ChengchenZhao/DrSeq2 |
| EMEP | nmf | 10.1093/bioinformatics/bty1056 | lixt314/EMEP |
| ESAT | pca,ica | 10.1101/gr.207902.116 | garber-lab/ESAT |
| FastProject | pca,fastica,kpca,t-sne,isomap,mds,spectral | N/A | YosefLab/FastProject |
| FiRE | low-dimensional sketch | 10.1038/s41467-018-07234-6 | princethewinner/FiRE |
| flotilla | pca,ica,tsne | N/A | yeolab/flotilla |
| GenePattern | pca | 10.12688/f1000research.15830.1 | genepattern/single_cell_clustering_notebook |
| gficf | pca,lsa,tsne,umap | 10.3389/fgene.2019.00734 | dibbelab/gficf |
| GiniClust | pca,tsne | N/A | lanjiangboston/GiniClust |
| GiniClust3 | pca,umap | 10.1101/788554 | rdong08/GiniClust3 |
| Granatum | correlation tsne,pca | 10.1186/s13073-017-0492-3 | lanagarmire/Granatum |
| GrandPrix | gaussian process mixture model | 10.1101/227843 | ManchesterBioinference/GrandPrix |
| Harmony | pca,user-supplied | 10.1101/461954 | immunogenomics/harmony |
| I-Impute | pca | 10.1101/772723 | xikanfeng2/I-Impute |
| IA-SVA | pca,ica,svd | 10.1038/s41598-018-35365-9 | UcarLab/iasva |
| iS-CellR | pca | 10.1093/bioinformatics/bty517 | immcore/iS-CellR |
| ivis | siamese neural networks | 10.1038/s41598-019-45301-0 | beringresearch/ivis |
| Jackstraw | pca | 10.1093/bioinformatics/btu674 | ncchung/jackstraw |
| LandSCENT | diffusion maps | N/A | ChenWeiyan/LandSCENT |

| | | | |
|---|---|---|---|
| LIGER | tsne,umap,pca,ica,nmf | 10.1016/j.cell.2019.05.006 | MacoskoLab/liger |
| MAGIC | pca | 10.1016/j.cell.2018.05.061 | KrishnaswamyLab/MAGIC |
| MELD | pca,phate | 10.1101/532846 | KrishnaswamyLab/MELD |
| Moana | pca | 10.1101/456129 | yanailab/moana |
| MOFA | pca | 10.15252/msb.20178124 | bioFAM/MOFA |
| Monocle | ddrtree | 10.1038/nbt.2859 | cole-trapnell-lab/monocle-release |
| m-pCMF | m-pcmf | 10.1101/496810 | pedrofale/mpCMF |
| MUDAN | pca,lda | N/A | JEFworks/MUDAN |
| NetNMF-sc | netnmf-sc | 10.1101/544346 | raphael-group/netNMF-sc |
| netSmooth | pca,tsne | 10.1101/234021 | BIMSBbioinfo/netSmooth |
| net-SNE | tsne | 10.1016/j.cels.2018.05.017 | hhcho/netsne |
| Nimfa | various nmf methods | 10.1093/bioinformatics/btw607 | ccshao/nimfa |
| ODEGRfinder | nmf | 10.1101/543447 | hmatsu1226/ODEGRfinder |
| OnlinePCA | online pca methods | N/A | rikenbit/OnlinePCA.jl |
| pagoda2 | pca | N/A | hms-dbmi/pagoda2 |
| Palantir | pca | 10.1038/s41587-019-0068-4 | dpeerlab/Palantir |
| pCMF | pcmf | 10.1101/211938 | https://gitlab.inria.fr/gdurif/pCMF |
| Pcreode | diffusion maps | 10.1016/j.cels.2017.10.012 | cole-trapnell-lab/monocle-release |
| PHATE | phate | 10.1101/120378 | KrishnaswamyLab/PHATE |
| PIVOT | pca,tsne,diffusion map,mds,penalizedlda | 10.1186/s12859-017-1994-0 | qinzhu/PIVOT |
| PyMINEr | N/A | 10.1016/j.celrep.2019.01.063 | bitbucket.org/scottyler892/pyminer_release |
| RAFSIL | rf based | 10.1101/258699 | kostkalab.net/software.html |
| RayleighSelection | spectral-base | arxiv.org/abs/1811.03377 | CamaraLab/RayleighSelection |
| rCASC | pca | 10.1101/430967 | kendomaniac/rCASC |
| robustSingleCell | pca | 10.1101/543199 | asmagen/robustSingleCell |
| TRPCA | trpca | 10.1089/cmb.2018.0255 | macieksk/rpca |
| RobustPCA* | rpca | 10.1145/1970392.1970395 | dganguli/robust-pca |
| sake | nmf | N/A | naikai/sake |
| SAUCIE | saucie | 10.1101/237065 | KrishnaswamyLab/SAUCIE |
| scAEspy | autoencoder | 10.1101/727867 | gitlab.com/cvejic-group/scaespy |
| scAlign | neural network | 10.1186/s13059-019-1766-4 | quon-titative-biology/scAlign |
| scanpy | pca,tsne,umap,diffusion map,draw graph | 10.1186/s13059-017-1382-0 | theislab/scanpy |
| scater | pca,mds,tsne,umap | 10.1093/bioinformatics/btw777 | davismcc/scater |
| scDeepCluster | denoising autoencoder | 10.1038/s42256-019-0037-0 | ttgump/scDeepCluster |
| Scedar | pca,umap,tsne | 10.1101/375196 | logstar/scedar |
| scell | pca | 10.1093/bioinformatics/btw201 | diazlab/SCell |
| scEpath | pca | 10.1093/bioinformatics/bty058 | sqjin/scEpath |
| scGEApp | tsne,umap,phate | 10.1101/544163 | jamesjcai/scGEApp |
| SCHNAPPs | pca,tsne,umap | N/A | C3BI-pasteur-fr/UTechSCB-SCHNAPPs |
| SCICAST | pca,svd,t-sne | N/A | C3BI-pasteur-fr/UTechSCB-SCHNAPPs |
| scMerge | pca,tsne | 10.1073/pnas.1820006116 | SydneyBioX/scMerge |

| | | | |
|---|---|---|---|
| scMetric | tsne | 10.1101/456814 | XuegongLab/scMetric |
| scNBMF | scnbmf | 10.1186/s12918-019-0699-6 | sqsun/scNBMF |
| SCORPIUS | lmds | 10.1101/079509 | rcannood/SCORPIUS |
| scPanoView | pca,tsne | 10.1371/journal.pcbi.1007040 | mhu10/scPanoView |
| scPred | pca | 10.1101/369538 | IMB-Computational-Genomics-Lab/scPred |
| scRCMF | nmf | 10.1109/TBME.2019.2937228 | XiaoyingZheng121/scRCMF |
| SCRL | scrl | 10.1093/nar/gkx750 | SuntreeLi/SCRL |
| scRMD | pca | 10.1101/459404 | ChongC1990/scRMD |
| scSeqR | pca,umap,tsne,diffusion map | N/A | rezakj/scSeqR |
| scSVA | tsne,pca,diffusion maps,umap | 10.1101/512582 | klarman-cell-observatory/scSVA |
| scUnifrac | pca,tsne | 10.1101/333393 | liuqivandy/scUnifrac |
| scVI | scvi | 10.1038/s41592-018-0229-2 | YosefLab/scVI |
| scvis | scvis | 10.1038/s41467-018-04368-5 | bitbucket.org/jerry00/scvis-dev |
| seurat | pca,ica,tsne,umap,phate,diffusion map | 10.1038/nbt.3192 | satijalab/seurat |
| SHARP | rp | 10.1101/461640 | shibiaowan/SHARP |
| simlr | pca,tsne | 10.1038/nmeth.4207 | BatzoglouLabSU/SIMLR |
| sincell | pca,ica,tsne,mds | 10.1093/bioinformatics/btv368 | RausellLab/Sincell |
| singleCellTK | pca,tsne,umap | 10.1101/329755 | compbiomed/singleCellTK |
| singlet | sam | N/A | iosonofabio/singlet |
| SinNLRR | nmf | 10.1093/bioinformatics/btz139 | zrq0123/SinNLRR |
| slalom | pca | 10.1186/s13059-017-1334-8 | bioFAM/slalom |
| Sleepwalk | N/A | 10.1101/603589 | anders-biostat/sleepwalk |
| slingshot | pca,diffusion map | 10.1186/s12864-018-4772-0 | kstreet13/slingshot |
| SoptSC | pca,nmf,tsne | 10.1101/168922 | WangShuxiong/SoptSC |
| spade | N/A | 10.1038/nbt.1991 | nolanlab/spade |
| SPRING | pca,spring | 10.1101/090332 | AllonKleinLab/SPRING |
| sscClust | pca,tsne,umap | 10.1016/j.gpb.2018.10.003 | Japrin/sscClust |
| SWNE | nmf | 10.1016/j.cels.2018.10.015 | yanwu2014/swne |
| TCM | N/A | 10.1038/s41467-018-05112-9 | gongx030/tcm |
| TSEE | elastic embedding | 10.1186/s12864-019-5477-8 | ShaokunAn/TSEE |
| UNCURL | pca,tsne,mds | 10.1093/bioinformatics/bty293 | yjzhang/uncurl_python |
| URD | pca,tsne,diffusion map | 10.1126/science.aar3131 | farrellja/URD |
| VAMF | vamf | 10.1101/166736 | willtownes/vamf |
| VASC | variational autoencoder | 10.1016/j.gpb.2018.08.003 | wang-research/VASC |
| VISION | pca,user-supplied | 10.1038/s41467-019-12235-0 | YosefLab/VISION |
| VPAC | vpac | 10.1101/523993 | ShengquanChen/VPAC |
| ZIFA | zifa | 10.1186/s13059-015-0805-z | epierson9/ZIFA |
| ZINB-WaVE | zinb-wave | 10.1038/s41467-017-02554-5 | drisso/zinbwave |
