## Supplemental Figures for "Supervised Application of Internal Validation Measures to Benchmark Dimensionality Reduction Methods in scRNA-seq Data"

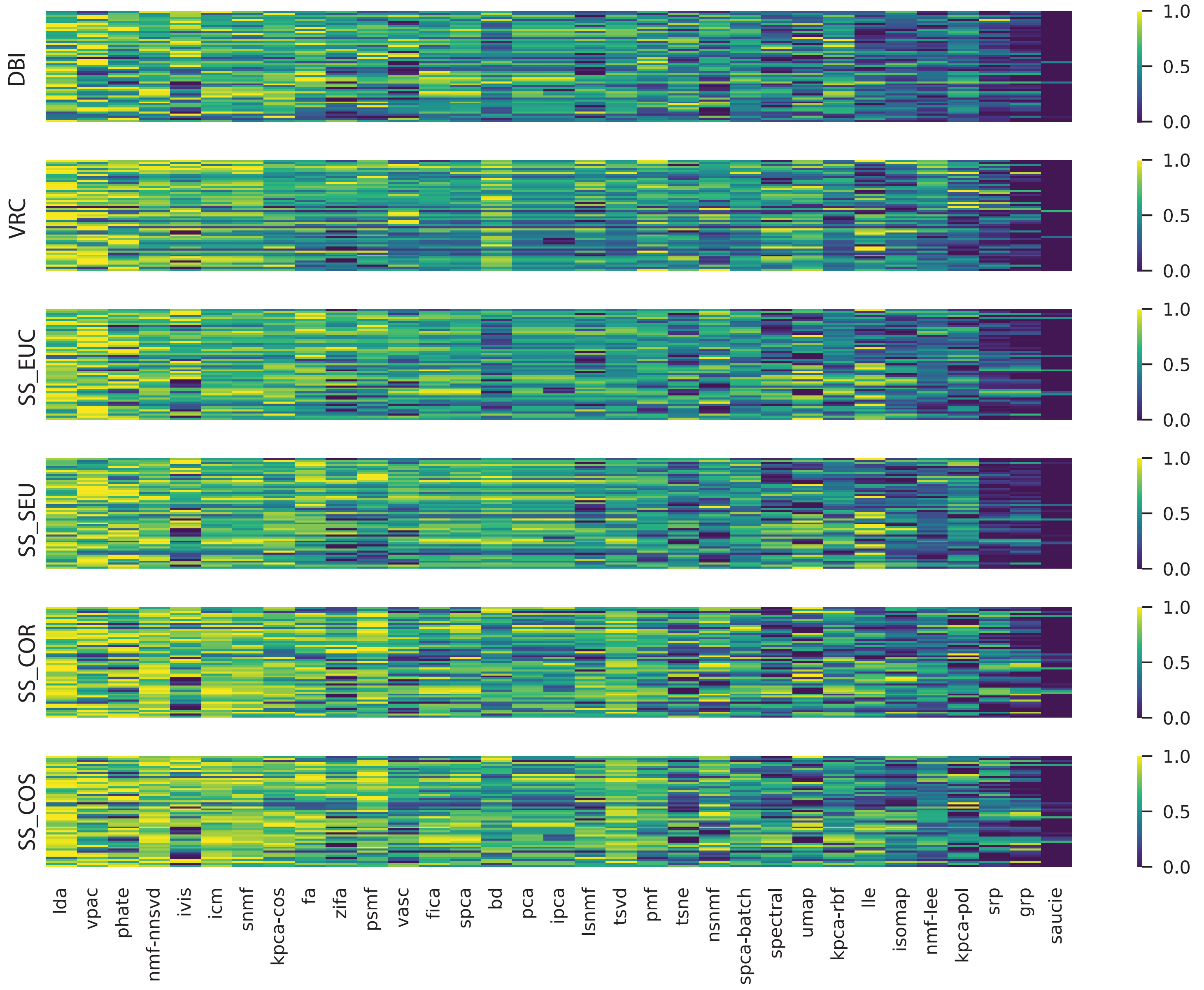


**Supplementary Fig. 1**: Heatmap of the 3 IVMs (SS, VRC and DBI) for each dimensionality reduction method (columns). Values were standardized using min-max normalization applied per-dataset (rows) to highlight differences between methods. SS measures with four distance metrics namely Euclidean (EUC), correlation (COR), cosine (COS) and standardized Euclidean (SEU) were shown.

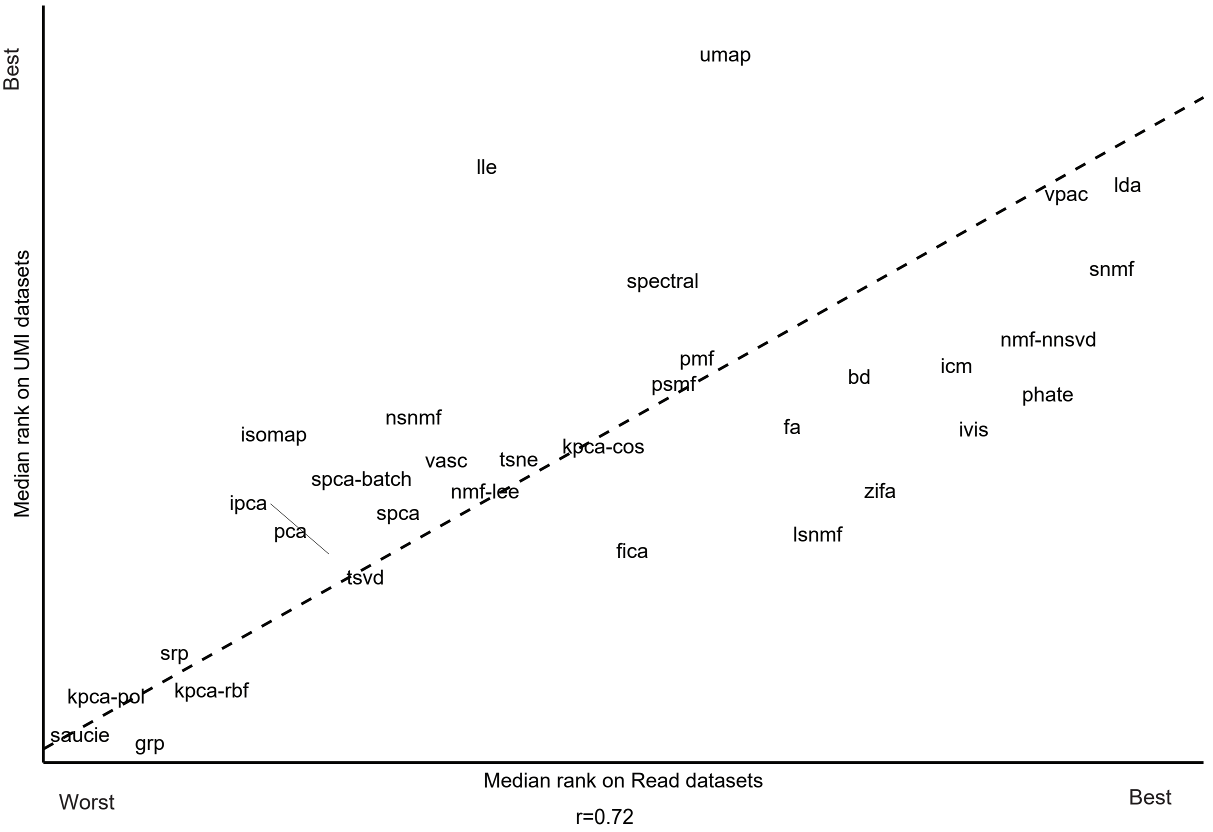


**Supplementary Fig. 2**: Scatterplot of method performance across UMI- and read- datasets. based Dotted line: parity line (y=x). r: Pearson correlation coefficient.

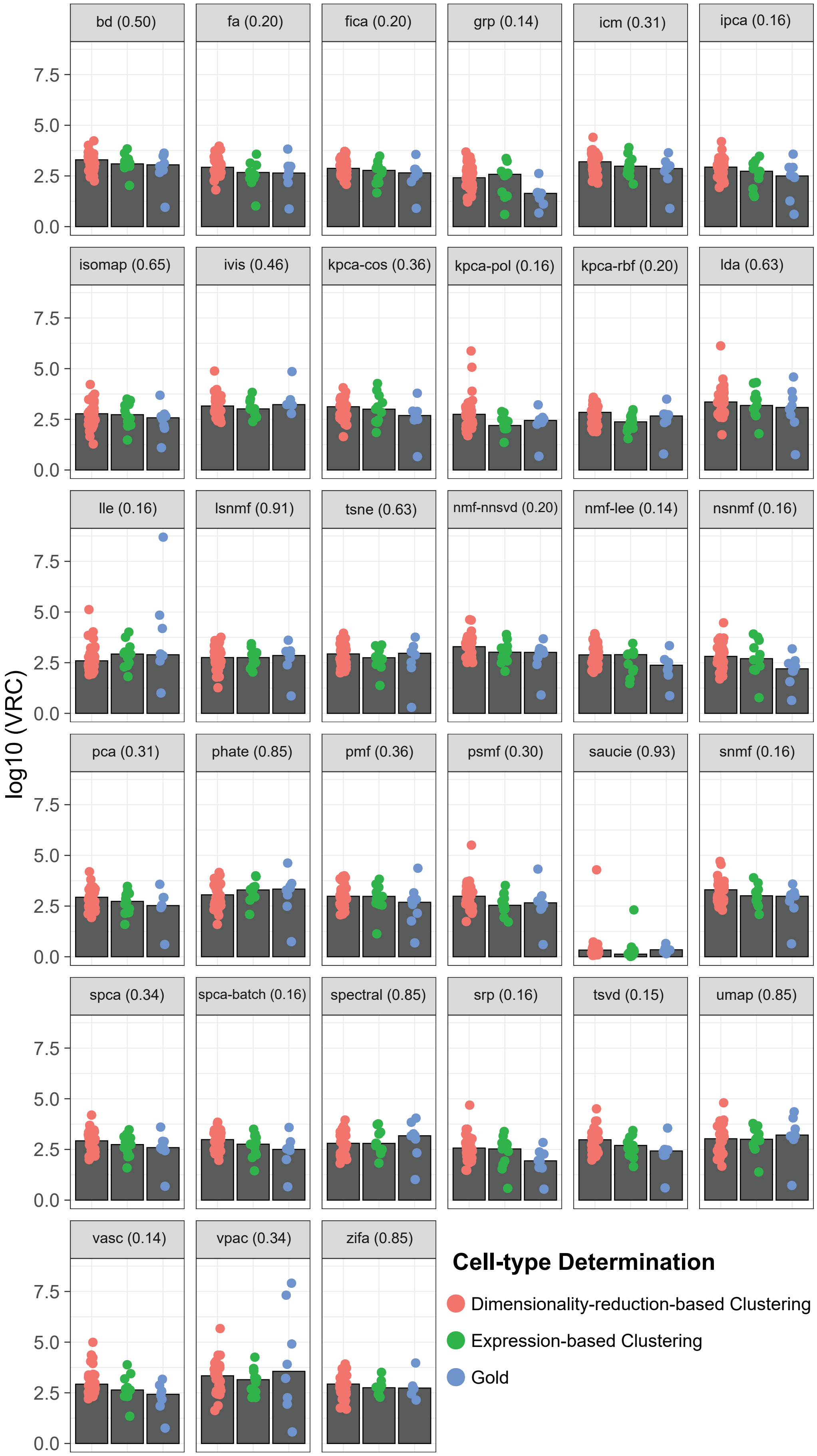


**Supplementary Fig. 3**: Comparison of Variance Ratio Criterion (VRC) across different cell-type annotation methods. Number in () shows FDR corrected ANOVA p-values comparing three groups. Datasets were classified according to the method the original publication used to determine their cell-type annotations. See Supplementary Table 1 for details on dataset classification. Gold: datasets where cell-type labels derive from a gold-standard truth (n=8). Expression-based clustering: datasets in which clustering was performed on gene expression to determine cell-type labels (n=11). Dimensionality-reduced-based Clustering: datasets in which dimensionality reduction was performed prior to clustering for determining cell-type labels (n=36).

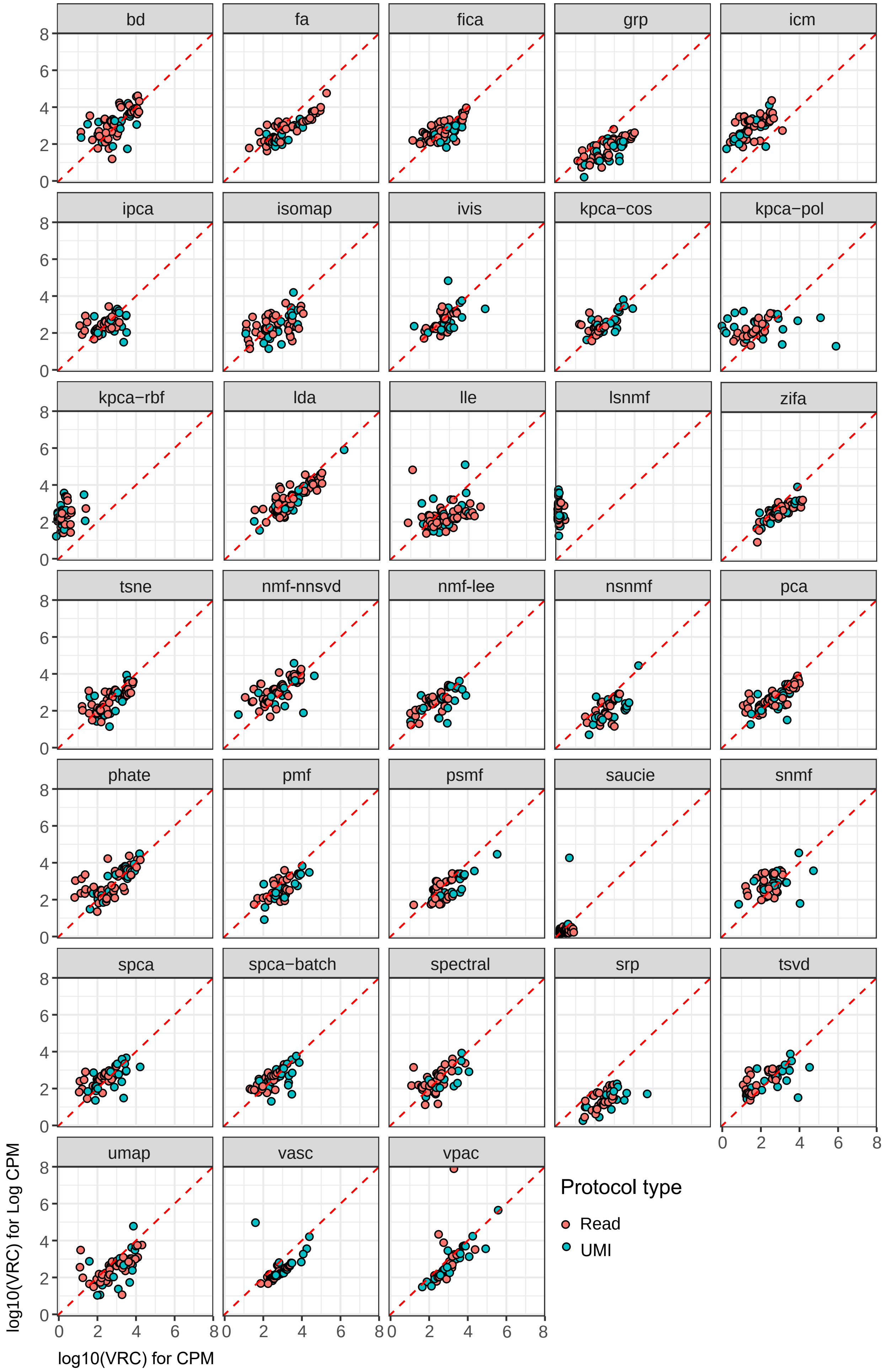


**Supplementary Figure 4:** Scatterplots of the effect of log transformation on Variance Ratio Criterion (VRC). Each point represents a dataset. Dotted red line: y = x. CPM: counts per million. A point positioned above the line indicates that the method produced a higher VRC for that dataset when applied to log(CPM+1) values rather than CPM

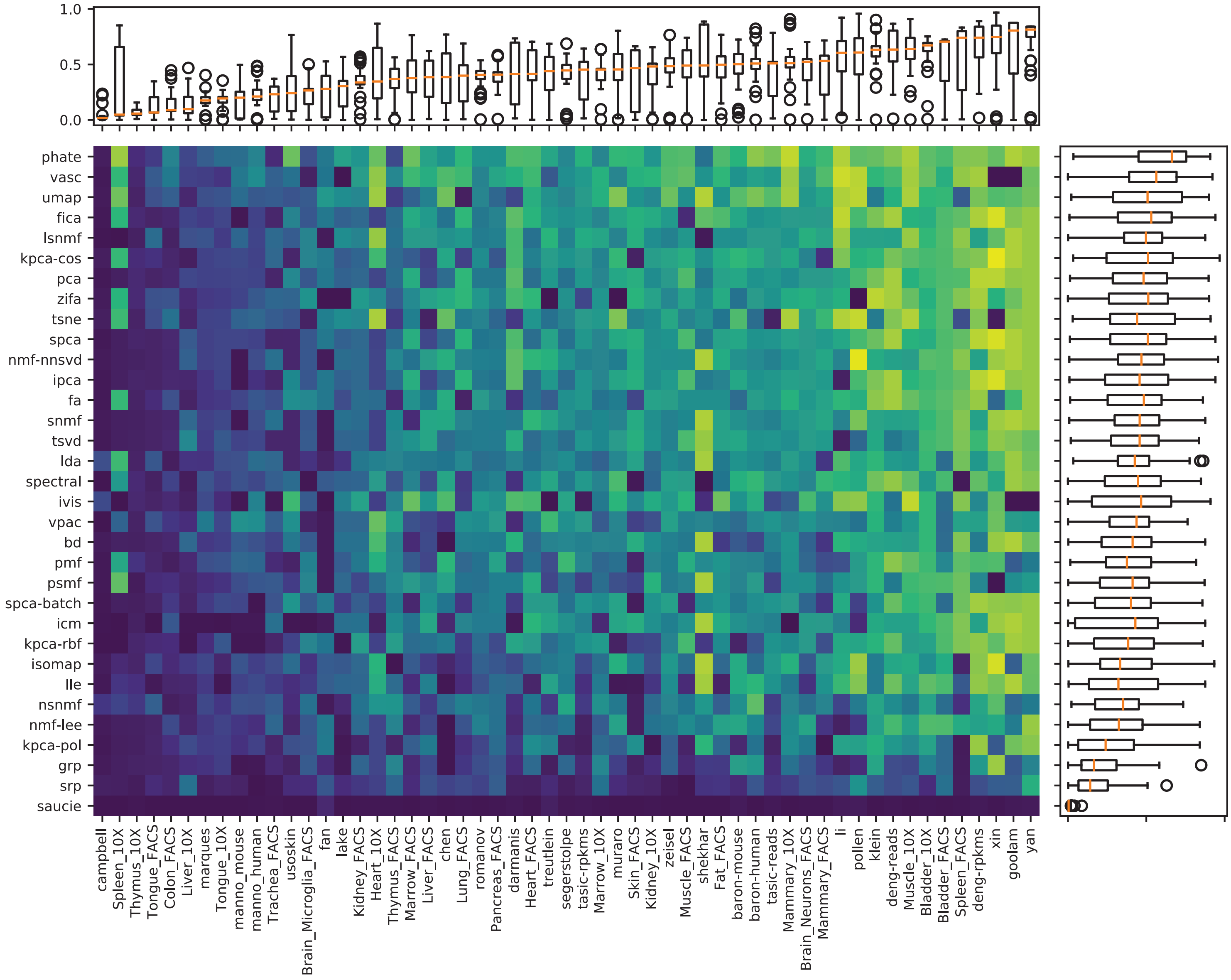


**Supplementary Fig. 5**: Heatmap with boxplots of the Normalized Mutual Information (NMI) achieved by each method (rows) for each dataset (columns).

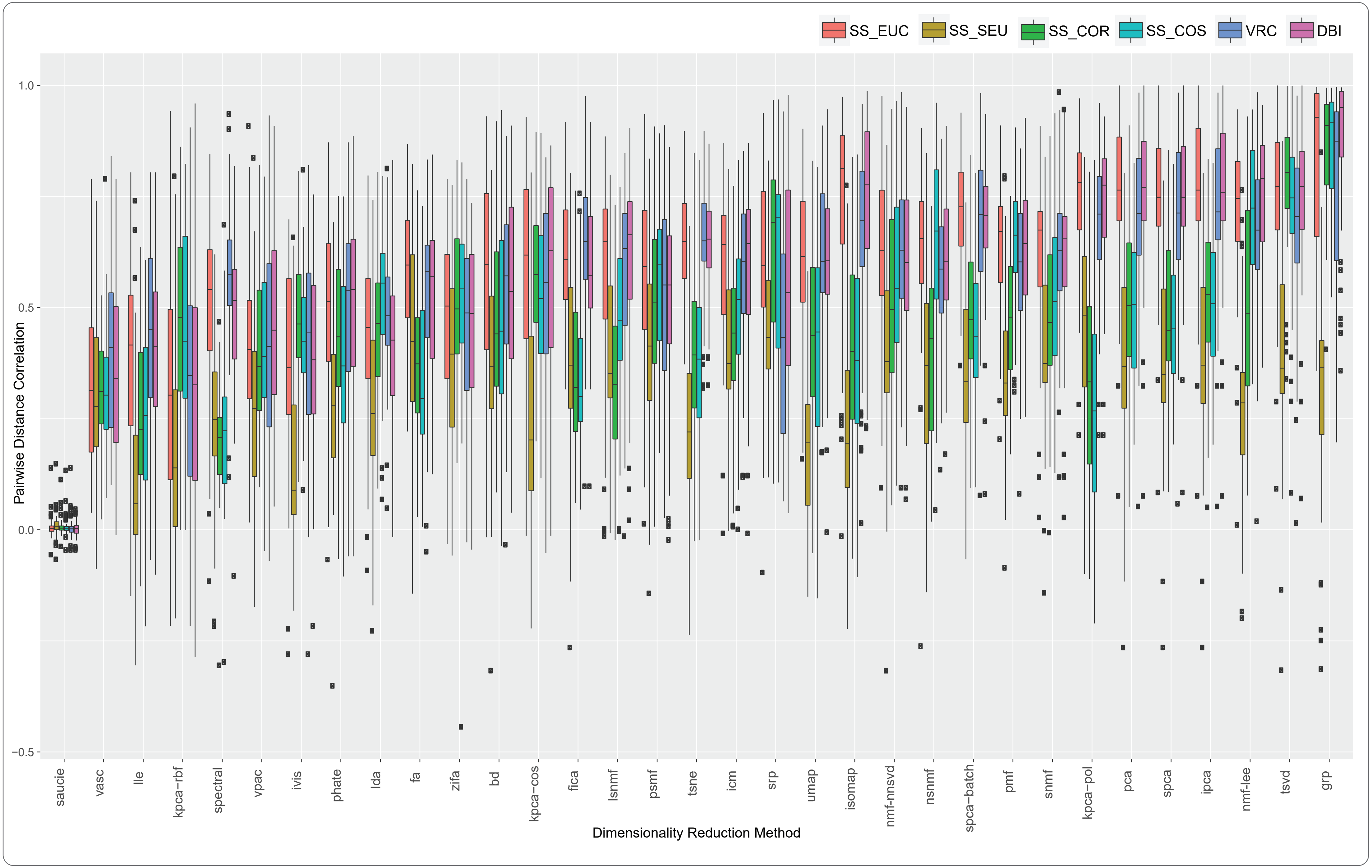


**Supplementary Fig. 6:** Effect of dimensionality reduction on global structure. For each full-dataset and low-dimension embedding, pairwise distances were calculated for the following similarity measures: Euclidean (EUC), standardized Euclidean (SEU), correlation (COR), and cosine (COS). To test whether global structure was well preserved, Spearman correlations were calculated between the full-dataset and low-dimension embedding, the idea being that the relative distance between points should not change after dimensionality reduction if global structure is preserved.

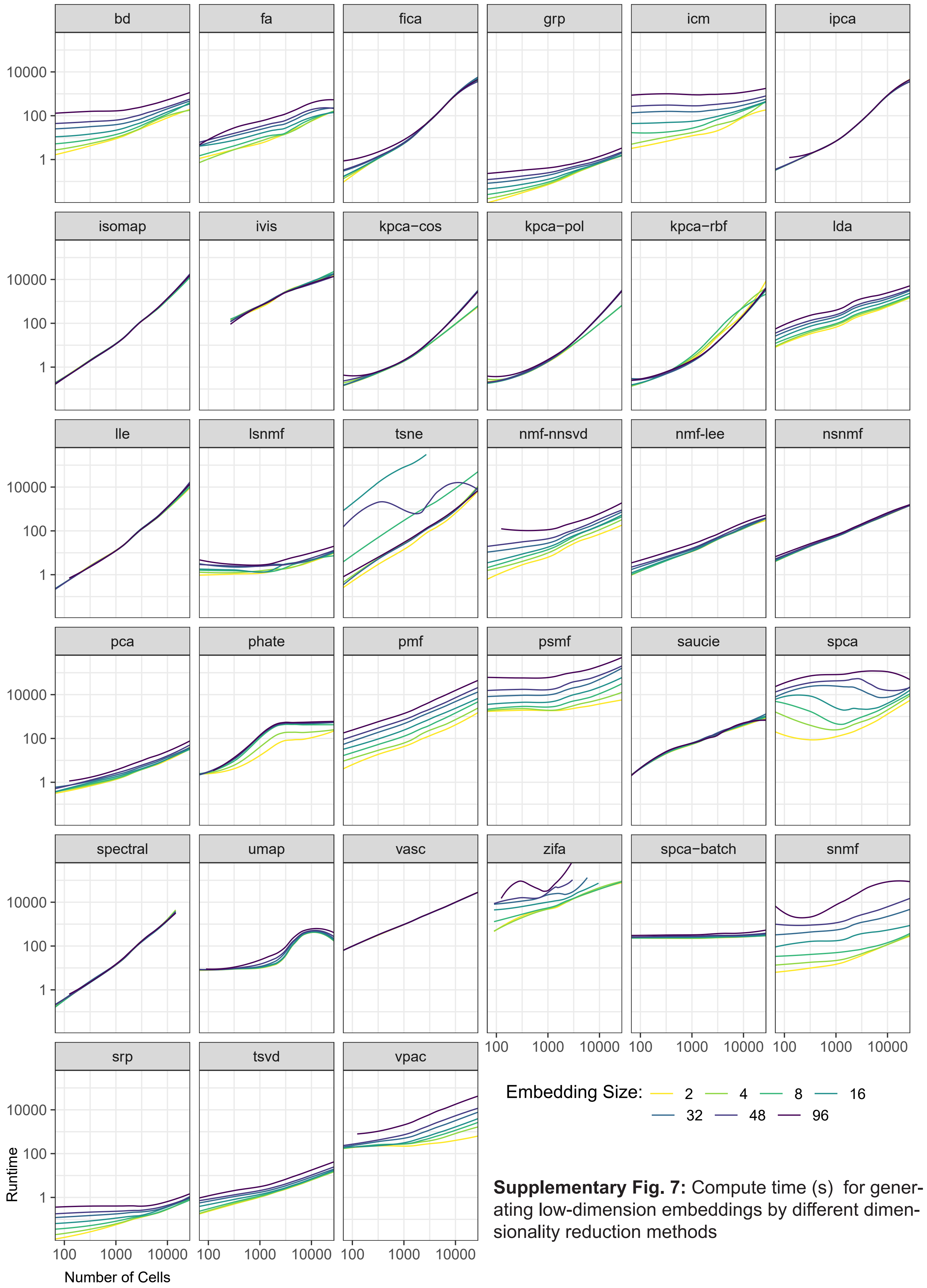


**Supplementary Fig. 7:** Compute time (s) for generating low-dimension embeddings by different dimensionality reduction methods

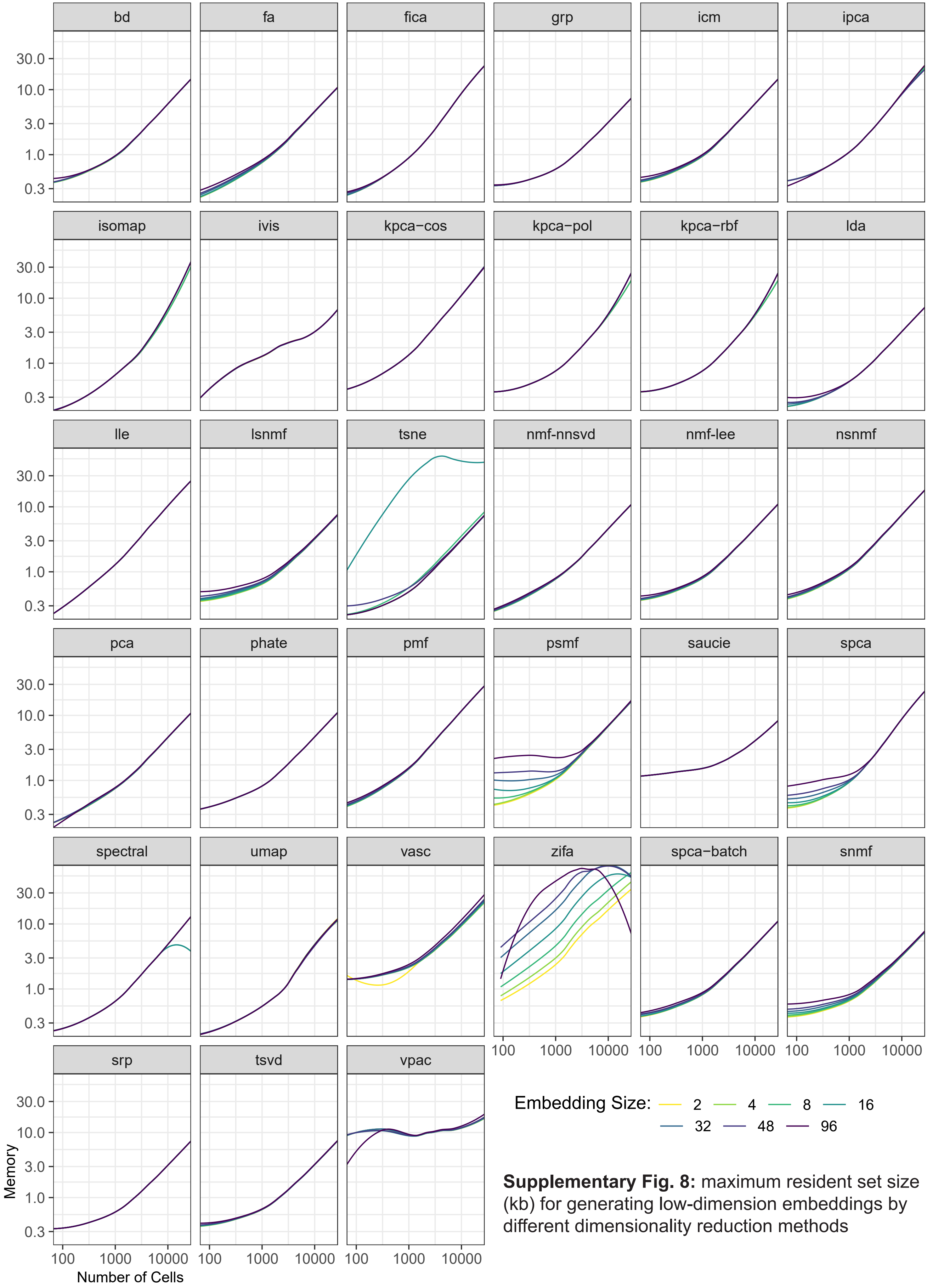


**Supplementary Fig. 8:** maximum resident set size (kb) for generating low-dimension embeddings by different dimensionality reduction methods
